# microRNAs as Indicators into the Causes and Consequences of Whole Genome Duplication Events

**DOI:** 10.1101/2021.09.01.458616

**Authors:** Kevin J. Peterson, Alan Beavan, Peter Chabot, Mark L. McPeek, Davide Pisani, Bastian Fromm, Oleg Simakov

## Abstract

Whole genome duplications (WGDs) have long been considered the causal mechanism underlying the dramatic increase in vertebrate morphological complexity relative to invertebrates. This is due to the retention and neo-functionalization of paralogues generated during these events, evolving new regulatory circuits, and ultimately morphological novelty. Nonetheless, an alternative hypothesis suggests that behind the retention of most paralogues is not neo-functionalization, but instead the degree of the inter-connectivity of the intended gene product, as well as the mode of the WGD itself. Here, we explore both the causes and consequences of WGD by examining the distribution, expression, and molecular evolution of microRNAs (miRNAs) in both gnathostome vertebrates as well as chelicerate arthropods. We find that although the number of miRNA paralogues tracks the number of WGDs experienced within the lineage, few of these paralogues experienced changes to the seed sequence, and thus are functionally equivalent relative to their mRNA targets. Nonetheless, the paralogues generated by the gnathostome 2R allotetraploidization event are retained in higher numbers on one sub-genome relative the second, with the miRNAs found on the preferred set of paralogons showing both higher expression of mature miRNA transcripts and slower molecular evolution of the precursor miRNA sequences. Importantly, WGDs do not result in the creation of miRNA novelty, nor do WGDs correlate to increases in complexity. Instead, it is the number of miRNA seed sequences in the genome itself that not only better correlate to instances in complexification, but also mechanistically explain why complexity increases when new miRNA families are established.

---

The origins of vertebrate complexity relative to most invertebrate taxa have long been sought in whole genome duplication (WGD) events (Ohno 1970). Various vertebrate lineages have experienced WGDs, with one (known as 1R) occurring after the divergence of the vertebrate lineage from invertebrates, but before the vertebrate last common ancestor (LCA), and a second (known as 2R) after the divergence of gnathostomes from cyclostomes, but before the gnathostome LCA (Hokamp et al. 2003; Lunin et al. 2003; Dehal & Boore 2005; Putnam et al. 2008; Simakov et al. 2020; Lamb 2021; Nakatini et al. 2021). Each of these two rounds of WGD would have doubled the genic content of the organism, and although most of these newly duplicated genes would be lost, some – through what Ohno (1970) called “forbidden mutations” – would be retained and now able to explore new evolutionary avenues normally not available to the gene product. Through this process of neofunctionalization, these genes would find new roles to play in vertebrate biology, and as Ohno (1970) first argued, would ultimately allow for an increase to their organismal complexity relative to most invertebrates (see also Holland et al. 1994; Sidow 1996; Escriva et al. 2006; Freeling and Thomas 2006; Putnam et al. 2008; Van de Peer et al. 2009; Huminiecki and Heldin 2010; Canestro et al. 2013; Glasauer and Neuhauss 2014; Van de Peer et al. 2017; Yamada et al. 2021).

Nonetheless, as Ohno (1970) also recognized, an alternative explanation behind gene retention following WGDs could be for reasons that have nothing to do with genic novelty *per se*. The gene-dosage (or gene-balance) model of selective gene retention (Birchler et al. 2001; Veitia 2002; Papp et al. 2003; Freeling and Thomas 2006) posits that genes whose products interact with other gene products in precisely determined stoichiometric ratios – in particular genes that encode for transcription factors, components of signal transduction pathways, and cell-cycle proteins – are selectively retained following WGDs, in contrast to gene products that are not under similar constraints, and hence return to single copy genes following WGDs (see also Blanc and Wolfe 2004; Seoighe and Gehring 2004; Birchler et al. 2005; Blomme et al. 2006; Conant and Wolfe 2008; Edger and Pires 2009; Hufton et al. 2009; Kassahn et al. 2009; Makino et al. 2009; Huminiecki and Heldin 2010; Makino and McLysaght 2010; Birchler and Veitia 2012; Buggs et al. 2012). Thus, the loss of newly generated paralogues from WGDs is not random with respect to the encoded gene product, but instead dependent upon its connectivity to other gene products (Gibson and Spring 1998; Veron et al. 2007). Consistent with this insight, dosage-sensitive genes – at least in human – are rarely found in tandem pairs, are often associated with haploinsufficiency, have significantly more protein interactions than the genomic mean, and are enriched in collections of disease-related genes relative to dosage-insensitive genes (Kondrashov and Koonin 2004; Birchler et al. 2005; Blomme et al. 2006; Makino and McLysaght 2010; Birchler and Veitia 2012; Singh et al. 2012).

Beyond the functional categorization of the gene product, a second reason why the loss of paralogues following WGDs is often not random involves the mode of the WGD event itself. There are two types of WGD (Ohno 1970; Garsmeur et al. 2014). The first – autopolyploidy – is when a mistake in DNA replication occurs relative to cytokinesis (Comai 2005) generating an entire second copy of the organism’s genome. Because of this identity, the subsequent elimination of these newly generated paralogues during the re-diploidization process is effectively random with respect to which of the two genomes housed the newly lost gene (Garsmeur et al. 2014). However, in instances of allopolyploidy in which two different diploid species hybridize, bringing together two distinct genomes into a single cell, the subsequent loss of paralogues – known as *homeologues* to distinguish them from the *ohnologues* generated during autotetraploidy (Glover et al. 2016) – is decidedly non-random. Instead, this rediploidization process results in “sub-genome dominance” or “genome fractionation” where one of the two hybridized genomes is preferentially retained relative to the other (Thomas et al. 2006; Schnable et al. 2011; Garsmeur et al. 2014; Session et al. 2016; Cheng et al. 2018; Edger et al. 2018). Therefore, during instances of autotetraploidy, biases in gene retention will be seen with specific kinds of genes in terms of their encoded gene products, but in instances of allotetraploidy, biases in gene retention will be seen both with respect to the kind *and* the genomic location of the gene itself for reasons that have nothing to do with potential neo-functionalizations.

Because of the non-randomness of paralogue losses from one of the two genomes following a hybridization event, allotetraploidy can be readily discerned from autotetraploidy simply by demonstrating the biased retention of genes from one sub-genome relative to the other (Session et al. 2016). Simakov et al. (2020) explored retention rates of paralogues across select vertebrate genomes and discovered that 1R was an autotetraploidy event (Fig. 1A), recognized by the parity of gene retention between sub-genomes “1” and “2” for both dosage insensitive (e.g., DNA repair proteins,Fig. 1B, left) as well as dosage sensitive (e.g., transcription factors, Fig. 1B, right) gene products (see Supp. Tables 1 and 2, and Supp. File 1). However, 2R was an allotetraploidy event where two different species – termed α and β by Simakov et al. (2020) – hybridized (see also Nakatani et al. 2021). Losses then preferentially accrued on the DNA derived from β paralogons relative to α paralogons, again for both dosage-insensitive and dosage-sensitive gene products (Fig. 1B).

**Figure 1.**
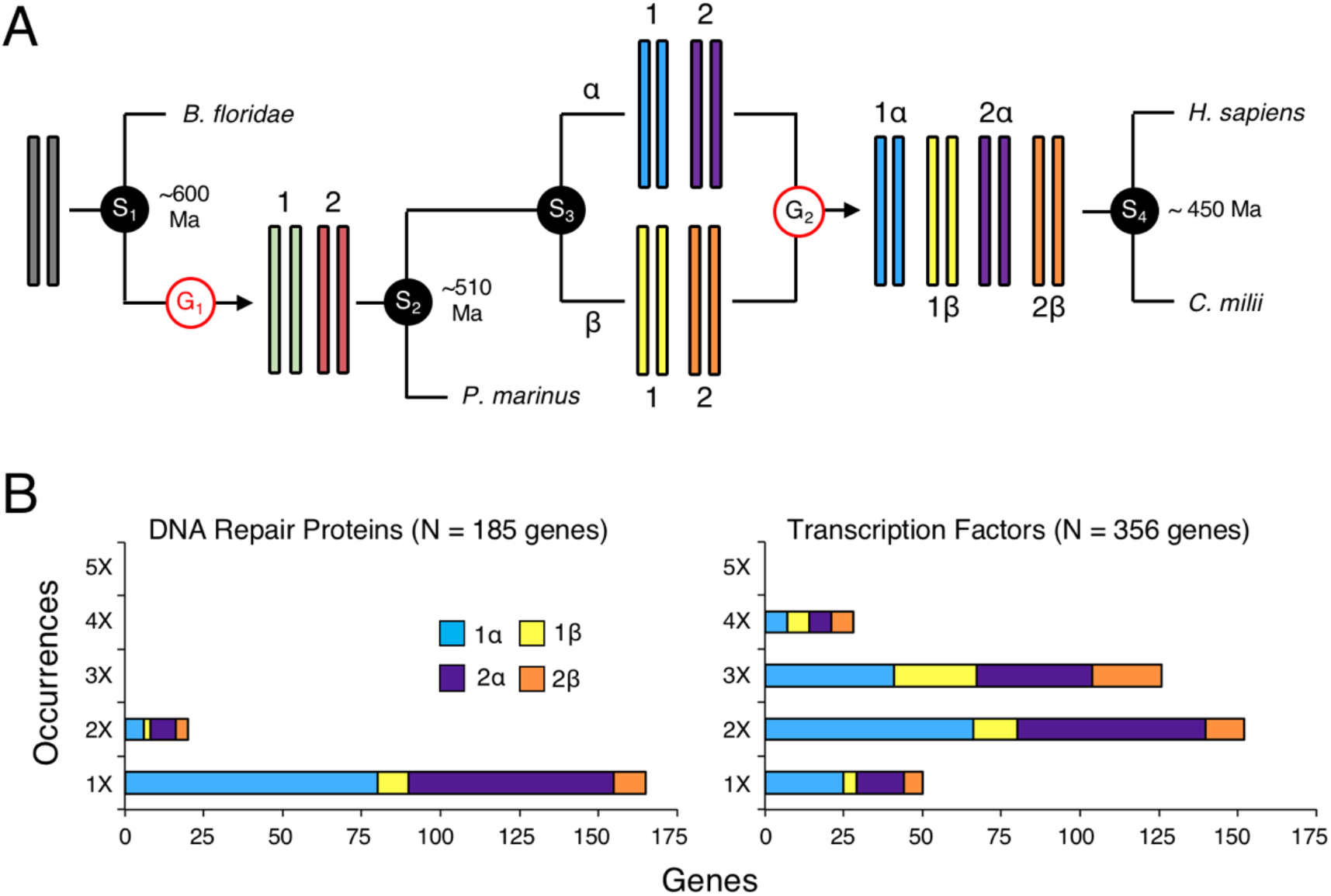
The Simakov et al. (2020) model of vertebrate genome evolution. **A**. Starting from an initial diploid state of an early chordate ancestor, sometime after the split (speciation [S] event 1) from the invertebrate chordates (e.g., the amphioxus *Branchiostoma floridae*), but before the separation of the extant jawless fish (e.g., the lamprey *Petromyzon marinus*) from the jawed fish (S2), the genome doubled in content (G1) generating a tetraploid genome. Because the retention of genes does not differ between sub-genomes 1 and 2, Simakov et al. (2020) reconstructed this WGD as an autotetraploidy event. Then, sometime after the vertebrate LCA (S2), there was a speciation event (S3) generating two, now extinct, lineages that Simakov et al. (2020) delineated α and β. After this speciation event, but before the gnathostome LCA (S4), there was a hybridization event between two species, one belonging to each of these two lineages, resulting in an allotetraploidization event (G2). Thus, the gnathostome genome – represented by *Homo sapiens* and the elephant shark *Callorhinchus milii* – is now octoploid with respect to the ancestral chordate genome. **B**. Retention evidence for an auto-followed by an allotetraplodization event in the early vertebrate lineage. Shown are the genomic distributions of 175 genes that encode DNA repair proteins (left, updated from Wood et al. 2001) and 175 genes that encode transcription factors (right, Lambert et al. 2018) that were present as single copy genes in the chordate LCA (Supp. Tables 1 & 2; Supp. File 1). Each gene was placed on paralogon “1” or “2” and “α” or “β” following Simakov et al. (2020) and Lamb (2021), with each paralogue in the genome separated by more than 50 kb from any other paralogue. Importantly, even though transcription-factor encoding genes are maintained at a mean of 2X relative to the invertebrate amphioxus, whereas DNA-repair encoding genes are largely maintained as single copy, both show significant enrichment of genes on the α paralogons versus the β paralogons, but not between the 1 and 2 sub-genomes (Supp. Tables 1 & 2).

Why genes from one of the hybridized genomes is preferred over the other remains unknown. Several hypotheses have been proposed (reviewed in Edger et al. 2018). One idea focuses on the hypothesis that the interactions between gene products governs retention such that only partners derived from the same genome would be retained (e.g., Thomas et al. 2006; Veitia 2009; Buggs et al. 2012). A second idea is that the differential expression of genes governs retention such that genes are retained from the genome that generated the higher transcript abundance due to potentially epigenetic differences between the two sub-genomes (e.g., Gout et al. 2010; Session et al. 2016; Xu et al. 2019). We sought to discriminate between these two competing (but not necessarily mutually exclusive) hypotheses by examining the genomic distribution of microRNA (miRNA) genes across a representative sample of jawed vertebrates (gnathostomes), as well as other lineages that also experienced WGDs, in particular chelicerate arthropods. miRNAs encode ∼22 nt non-coding RNA products that interact with target messenger RNAs (mRNAs) primarily through nucleotide positions 2-8 of the mature miRNA gene product, what is known as the “seed” (Bartel 2009, 2018; Dexheimer & Cochella 2020). This interaction between the seed sequence of a miRNA and a target mRNA results in the abrogation of the mRNA through the activity of the protein Argonaute that forms the enzymatic core of the RNAi-induced silencing complex (Schirle et al. 2014; Nakanishi 2016). Because the interaction between the miRNA seed sequence and the target mRNA sequence involves simple base-pairing rules between the two, the same seed sequence from different miRNA paralogues can potentially interact with the same set of mRNA targets. This then allows a test between these two hypotheses for sub-genome dominance: if the selective retention of genes is primarily due to interactions between genic products – whether RNA or protein – this should result in randomness of miRNA retention between the α and β paralogons of extant gnathostomes, given the strong conservation of the seed and 3’-CR regions of gnathostome miRNAs (Fromm et al. 2015). Alternatively, if the reasons for sub-genome dominance center around the location of the gene itself, then miRNA paralogue retention should follow the same trends that Simakov et al. (2020) demonstrated for protein-coding genes (Fig. 1B).

Here, we show that similar to younger genome duplication events in fish (Berthelot et al. 2014) and *Xenopus* (Session et al. 2016) miRNAs follow the same retention trends as their principle targets and are selectively retained following WGDs. An examination of genomic retention unambiguously shows that gnathostome miRNAs – like their protein-coding genes – are selectively retained on the α genome relative to the β genome. However, unlike protein-coding genes (Simakov et al. 2020), miRNA paralogues are continually lost on the β paralogons relative to the α paralogons for hundreds of millions of years after the gnathostome LCA. Further, these β homeologues are expressed at lower levels, and experience more mutations to their mature sequence, than the homeologues found on α paralogons. Finally, following Heimberg et al. (2008), we argue that WGDs are not primary drivers of morphological evolution. Instead, the best predictor of morphological and behavioral complexity in any animal lineage is the number of distinct miRNA seed sequences present in the genome itself, sequences that, surprisingly, are not the result of WGDs.

## RESULTS

### Retention of microRNAs following WGD events

Aside from the studies of Bhambri et al. (2018) and Desvignes et al. (2021), most efforts to understand the increase in miRNA paralogue numbers in metazoan taxa that have undergone WGD events (e.g., Hertel et al. 2006; Berthelot et al. 2014; Braasch et al. 2016; Leite et al. 2016; Shingate et al. 2020a; Nong et al. 2021) were hampered by the difficulty in assigning direct homology between individual miRNA genes. However, MirGeneDB (Fromm et al. 2015, 2020) was created with the specific intent to use a consistent nomenclature system that explicitly recognizes paralogues within a taxon and orthologues across taxa based on both syntenic and phylogenetic analyses. Therefore, the miRNA repertoire of any one species can be directly compared to any other within the database. In addition, the miRNA repertoire of extinct species can be easily reconstructed given that the evolutionary point of origin of every miRNA within the database (nearly 15,000 robustly identified miRNA loci from 73 metazoan taxa), including its family-level membership, is explicitly identified within the context of the database’s taxonomy.

To assess our methodology with respect to miRNA homology assignments, we constructed a concatenated data set of all 254 precursor miRNA sequences reconstructed as present in the gnathostome LCA (Supp. File 2). Each of these 254 miRNA precursor sequences from 32 representative taxa was aligned, concatenated, and analyzed by Bayesian analysis (see Materials and Methods). Because the phylogeny of these 32 taxa is known, any deviation from this accepted topology could be due to one of several reasons including through mis-assignment of miRNA gene identities, or due to meiotic exchanges between homeologues that could have occurred after the hybridization event (Edger et al. 2018). However, we find robust support for this accepted topology with most nodes supported with high posterior probabilities (Supp.Fig. 1). The relatively low support of the eutherian nodes Glires and Atlanogenata is similarly difficult to capture with protein-coding genes (Supp.Fig. 1; see Materials and Methods), and thus appears to be clade-specific issues not related to difficulties with miRNA homology assignments.

Because our miRNA homology assignments appear robust, we next asked if the number of occurrences of miRNA paralogues corresponds to a taxon’s known number of genome duplication events (Fig. 2). Profiling the distribution of miRNA paralogues within the genome of the shark *C. millii* taxa shows that it has numerous instances of up to four paralogues of miRNAs (but no more) distributed throughout the genome with no paralogue separated by less than 50 kb from another, the distance used herein as the maximal extent of a miRNA polycistron (Baskerville and Bartel 2005). Interestingly, all of the occurrence of 2 or more occurrences of miRNA paralogues in the gnathostome genome involve miRNA families that evolved before the LCA of all living vertebrates (Fig. 2A, black bars), but none involving families that evolved after the vertebrate LCA (Fig. 2A, white bar). The teleost fish *D. rerio* though has up 7 paralogues of miRNAs due to the 3R event that occurred in the teleost lineage after it split from the holostean fish like the gar, but before the teleost LCA (Glasauer and Neuhauss 2014; Desvignes et al. 2021). For miRNA families that evolved after 2R, but before 3R, these occur at no more than two times in the genome of *D. rerio* (Fig. 2B, gray bars), and for those that evolved after 3R, these are found as genomic singletons (Fig. 2B, white bar).

**Figure 2.**
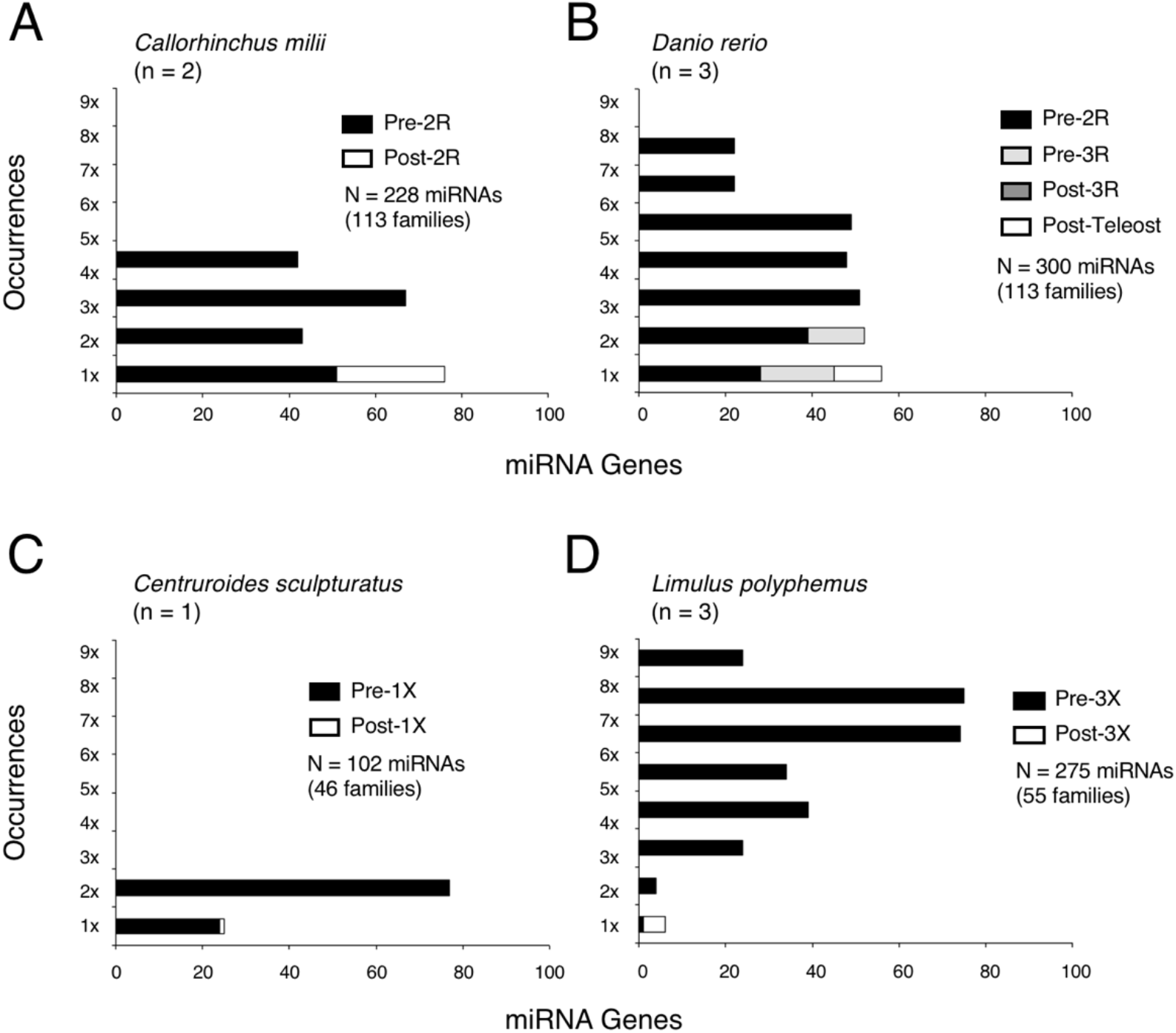
The occurrences of miRNA paralogues reflect the number of WGD events. Shown are the number of occurrences of miRNA paralogues in four different metazoan genomes, the gnathostomes *C. milii* (elephant shark) (**A**) and *D. rerio* (zebrafish) (**B**), and the chelicerates *C. sculpturatus* (bark scorpion) (**C**) and *L. polyphemus* (Atlantic horseshoe crab) (**D**). In each case the maximal number of miRNA paralogues separated by at least 50 kb is simply equal (or nearly equal) to 2^n^ where n is the number of WGDs. Importantly, none of these WGDs resulted in a significant increase in the number of miRNA families, only paralogues to previously existing families.

Similarly, within the chelicerates we find that again miRNAs track the number of WGD events. Most arthropods including the tick *I. scapularis* (Schwager et al. 2017; Shingate et al. 2020b) have not experienced any WGD, and thus have few if any miRNA paralogues separated by more than 50 kb from one another (MirGeneDB.org). However, the scorpion *C. sculpturatus*, which has experienced a single WGD shared with spiders (Schwager et al. 2017), has numerous miRNA paralogues on separate contigs, but none on more than two (Fig. 2C). Furthermore, the Atlantic horseshoe crab *L. polyphemus*, which like teleosts has undergone three WGDs (Nong et al. 2021), has miRNA paralogues occurring in the genome up to 8 times for miRNA families that evolved before the WGD events (Fig. 2D, black bars), but only singletons for miRNAs that evolved after the WGDs (Fig. 2D, white bar). Therefore, similar to the genes that encode a subset of their principle targets (Fig. 1B, right), miRNAs are retained as multiple paralogues following WGD events in both gnathostomes as well as in chelicerate arthropods, paralogue numbers that reflect the number of WGDs themselves.

### The Distribution of miRNAs in the Genomes of Three Last Common Ancestors

Because the gnathostome miRNAs are distributed throughout the genome in a manner that reflects the number of WGDs, we next sought to reconstruct the miRNA repertoire of three last common ancestors (Chordata, Vertebrata and Gnathostomata). Simakov et al. (2020) confirmed that the chordate LCA had at least 17 linkage groups (Putnam et al. 2008, Sacerdot et al. 2018; Lamb 2021; Nakatani et al. 2021), and related these ancestral linkage groups (ALG) to the extant genomes of four key chordate taxa – the amphioxus *B. floridae*, the chicken *G. gallus*, the spotted gar *L oculatus*, and the frog *X. tropicalis*. Thirty of 33 miRNA genes or gene clusters present in this LCA (MirGeneDB.org) could reliably be placed on one of these 17 ALGs (Supp. Fig 2A). Twenty-six of these miRNAs or clusters of miRNAs would be passed on to the vertebrate LCA, and all but one (Mir-33) are still found on the same ancestral ALG (Supp.Fig. 2A,B, pound sign); an additional four miRNAs or clusters of miRNAs were lost after the chordate LCA, but before the vertebrate LCA (Supp.Fig. 2A, downward arrows).

The vertebrate LCA is reconstructed as having 34 linkage groups (Supp.Fig. 2B), a result of the first WGD event with no apparent chromosomal fusions or fissions (Simakov et al. 2020; Lamb 2021; Nakatani et al. 2021). The three unplaced miRNA families in the chordate LCA (MIR-34, MIR-92, and MIR-103) are all now placed on the genomic reconstruction of the vertebrate LCA. Of the 32 miRNAs or clusters of miRNAs present in the Olfactores LCA, 17 are now present in two copies, one on each of the two sub-genomes, and thirteen on either sub-genome 1 or 2 (Table 1). The vertebrate LCA has an additional 43 miRNA genes or clusters of genes (Supp.Fig. 2B, bold) that evolved after the Olfactores LCA, but before the vertebrate LCA. Nineteen of these are found on both sub-genomes, and thus evolved before the autotetraploidy event. An additional 24 genes or clusters of genes are found on only one of the two sub-genomes (Supp.Fig. 2B, asterisks), either because one ohnologue was lost, or because it evolved sometime between 1R and the vertebrate LCA. Comparisons with the pre-vertebrate miRNAs indicates that the former is more likely as there is no statistical difference between the retention of pre-vertebrate singletons versus vertebrate singletons (χ2 = 1.50, *df*=1, *P*=0.22). Therefore, although these miRNA distribution data are best explained by an autotetraploidy event, 1R did not result in the evolution of an unusually high number of novel miRNA families (Heimberg et al. 2008).

**Table 1.**
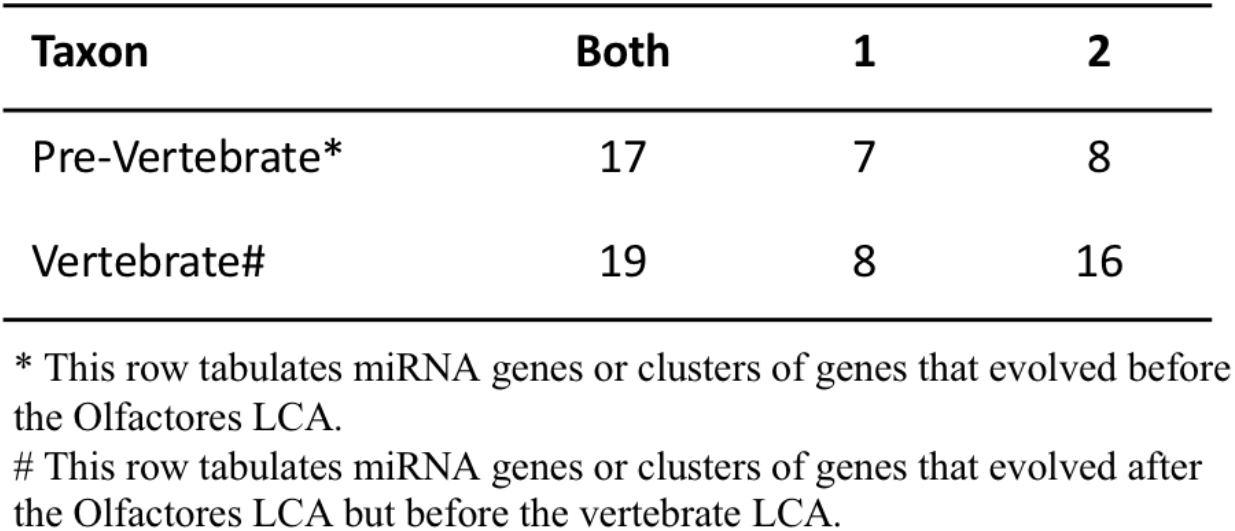
miRNA paralogue retention in each of the two original sub-genomes in the vertebrate LCA.

The genome of the gnathostome LCA is reconstructed as having at least 45 linkage groups, a result of the second tetraploidy accompanied with several chromosomal fusion events (Supp.Fig. 2C) (Simakov et al. 2020; Lamb 2021; Nakatani et al. 2021). Seventy-nine miRNA families were inherited from the vertebrate LCA, with paralogues distributed between one to four paralogons (Supp. File 3). An additional 11 families evolved after this second WGD, but before the gnathostome LCA (Supp.Fig. 2C, upward arrows), and are present as singletons in the genome of the gnathostome LCA. Thus, again consistent with Heimberg et al. (2008), 2R did not result in an influx of an unusually high number of new miRNA families into the gnathostome lineage (Fromm et al. 2015).

With this genomic reconstruction in hand, we can now ask if miRNAs show sub-genome dominance as a result of the 2R event, but not due to the 1R event.Figure 3 shows the paralogon distributions of the miRNAs in the genome of the gnathostome LCA, as well as in the genomes of five select descendant taxa. In all cases, miRNAs are significantly enriched on the α paralogons relative to the β paralogons (*χ*^*2*^ = 95.6, *df* = 1, *P*<0.0001), but are not enriched on sub-genome 1 relative to sub-genome 2. Thus, as demonstrated by Simakov et al. (2020) for protein-coding genes (Fig. 1), miRNAs follow the same genomic biases that resulted from 2R allotetraploidy event. However, unlike protein-coding genes (Simakov et al. 2020), miRNA losses continue on the β paralogons relative to the α paralogons long after the gnathostome LCA *(χ*^*2*^ = 32.0, *df* = 3, *P*<0.0001) (Table 2; Supp.Fig. 3). Thus, whatever the mechanism for biased gene retention following allotetraploidy events, this bias continues – at least for miRNAs – for hundreds of millions of years after the 2R event itself.

**Figure 3.**
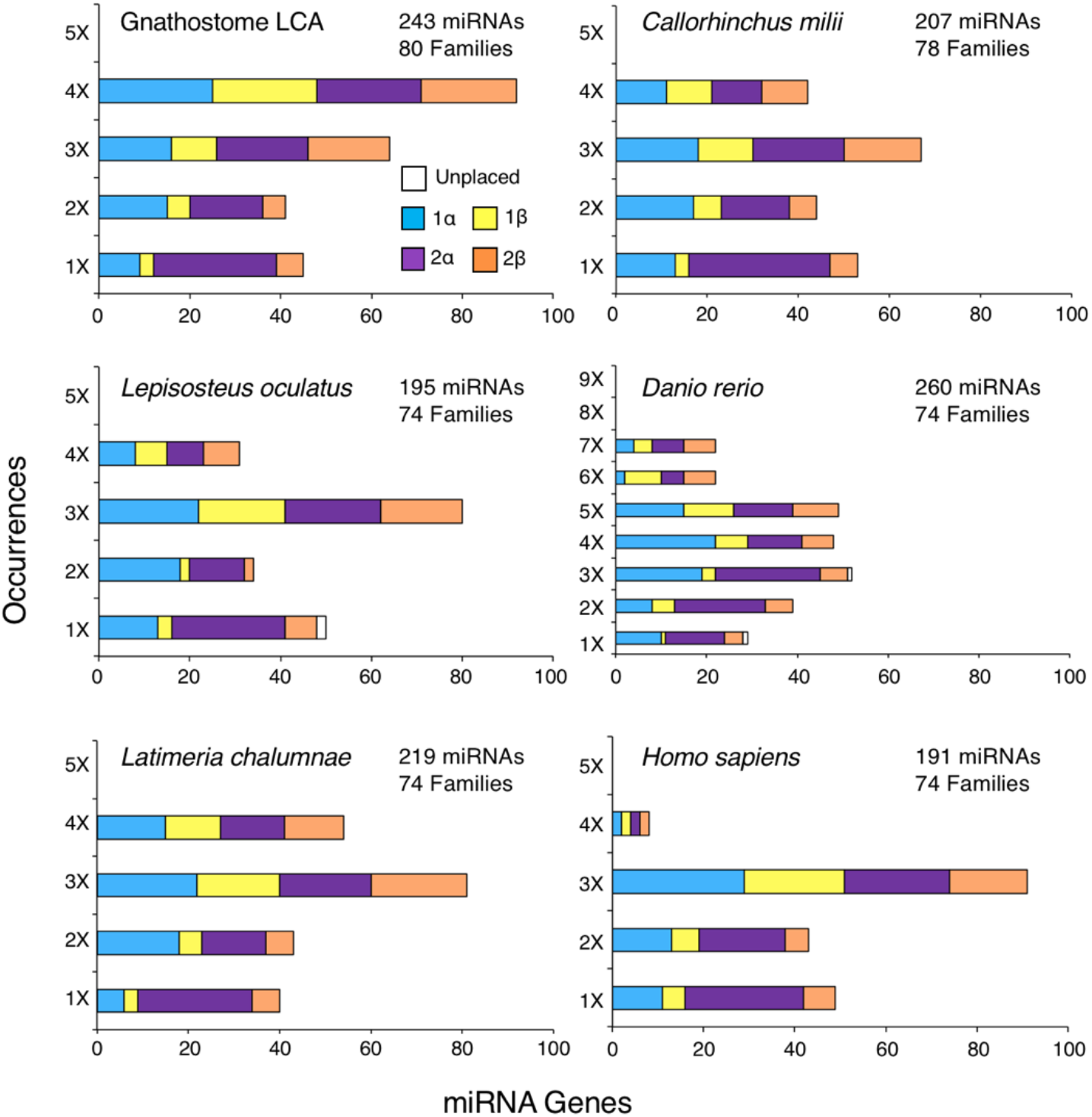
The distribution of miRNA paralogues across the four paralogons in six representative gnathostome taxa. Tabulating the occurrences of miRNAs paralogues of miRNA families present in the vertebrate LCA shows that in each instance these paralogues are enriched on α paralogons versus β paralogons, but not between 1 versus 2 sub-genomes (Table 2). These observations are consistent with Simakov et al.’s (2020) hypothesis that the gnathostome 2R events were an allotetraploidization following an autotetraploidization (see Fig. 1). Unexpectedly, in contrast to the protein-coding repertoire (Simakov et al. 2020), there is continued loss of β versus α paralogues long after the gnathostome LCA as seen in not only these five taxa, but in all extant gnathostome taxa analyzed (Table 2). Both the number of paralogues and the number of families remaining in each extant taxon in relation to the gnathostome LCA are also indicated.

**Table 2.** miRNA paralogue retention in each of the four ancestral paralogons in the gnathostome LCA.

|  | **1α** | **2α** | **1β** | **2β** |
| --- | --- | --- | --- | --- |
| Total | 65 | 86 | 41 | 50 |
| 4X* | 25 | 23 | 23 | 21 |
| 3X* | 16 | 20 | 10 | 18 |
| 2X* | 15 | 16 | 5 | 5 |
| 1X* | 9 | 27 | 3 | 6 |
| post-LCG# | 54 | 73 | 64 | 93 |

\* These rows tabulate the total number of miRNA genes present singularly or in clusters in four, three, two or one copies in the genome of the gnathostome LCA.

### miRNA Sequence Expression and Evolution

Because there is a clear distinction between the retention of miRNA genes on α versus β paralogons, and the mature sequences for each set of paralogues is functionally equivalent – at least with respect to the sequence of the seed and most of the 3’CRs (Fromm et al. 2015; Supp.Fig. 4) – we next asked if we could detect differences between either the expression of paralogon-specific sets of miRNAs, or the rate of primary sequence evolution of the pre-miRNAs themselves. We first compiled read data (standardized as reads per million) from MirGeneDB.org for 11 taxa where at least one α and one β paralogon houses a miRNA paralogue generated by one or both of the vertebrate WGD events (Supp. File 4). Importantly, only miRNAs with unique mature sequences were chosen for this analysis, greatly reducing the number of genes analyzed, but ensuring that the reads from multiple loci were not spuriously merged into one. Interestingly, the median expression from sub-genome 1 is half that of sub-genome 2, and from the β paralogons half that of α paralogons (Fig. 4A, C; Supp. Table 3) with the difference in median expression between α and β paralogons significant (Χ ^*2*^ = 63577.9, *df*=1, *P*<0.0001). Thus, as predicted from other independent allotetraploidy events (Session et al. 2016), the paralogons retaining the higher percentage of paralogues (in this case the α paralogons) also express miRNA genes at higher levels relative to the β paralogons. Unexpectedly though expression also shows a two-fold difference between the 1 versus 2 sub-genomes.

**Figure 4.**
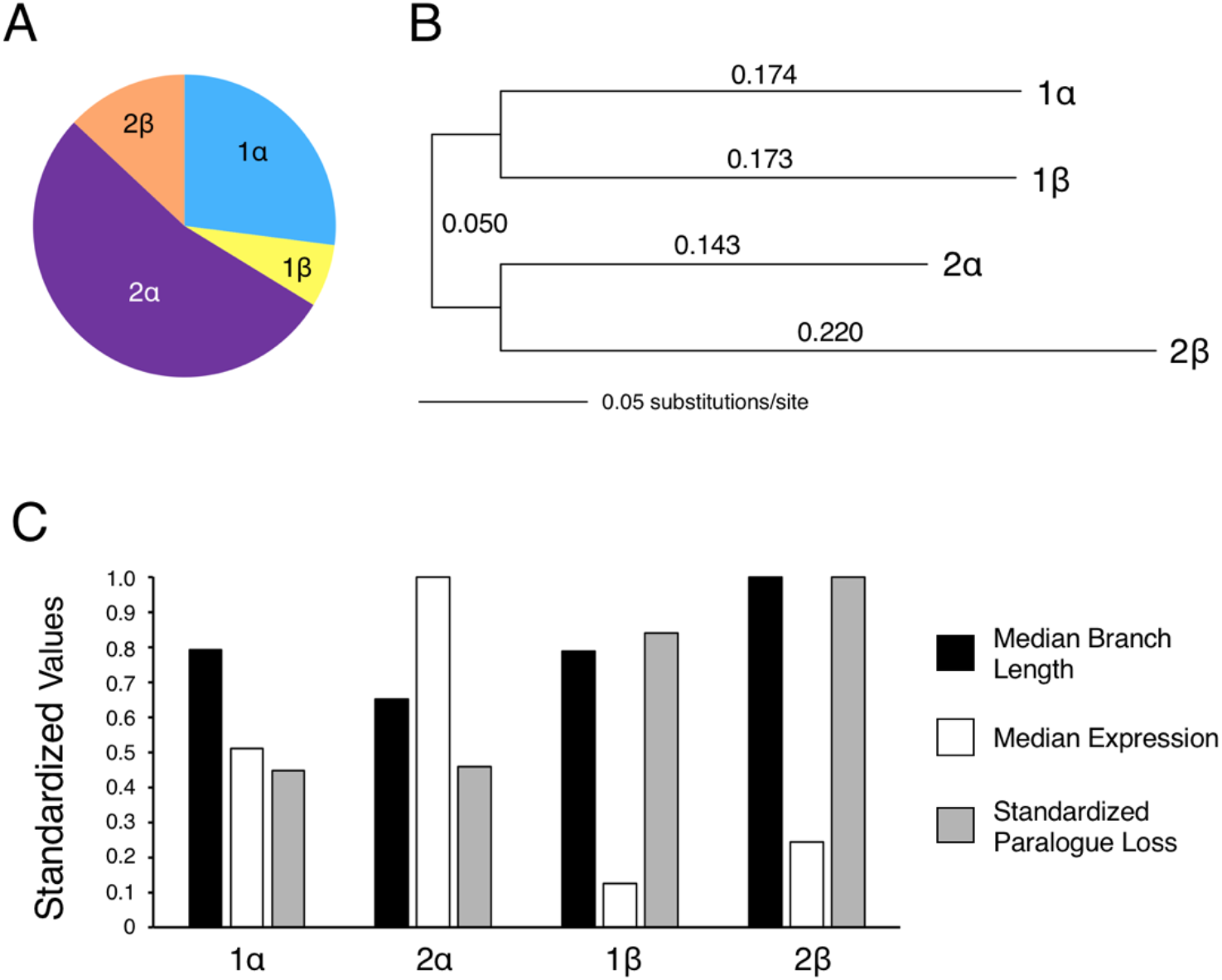
The expression and evolution of *α* versus *β* paralogues in extant gnathostomes. **A**. Tabulating the reads/million (rpm) values for unique miRNA mature sequences with at least one paralogue on an α paralogon and one on a β paralogon in 11 extant gnathostome taxa shows that expression is enhanced on α paralogons versus β paralogons (Supp. Table 4). **B**. Concatenating all paralogues present in 13 extant gnathostome taxa with at least one paralogue on an α paralogon and one on a β paralogon shows that pre-miRNAs present on the 2α paralogon evolved significantly slower, and those present on the 2β paralogon significantly faster, than the paralogues present on the sub-genome 1 paralogons (Supp. Table 5). This mid-point rooted phylogram was constructed using Maximum Likelihood (see Materials and Methods); branch lengths are as indicated. **C**. The inverse relationship between miRNA paralogue retention and pre-miRNA molecular evolution versus miRNA expression present across the four paralogons. The Y axis indicates the standardized values to maximum for median expression (rpm, Supp. Table 4), median branch lengths (Supp. Table 5), and standardized paralogue loss (Table 2). miRNA loci found on the α paralogons are expressed significantly higher and retain significantly more miRNA loci than miRNA loci found on β paralogons. Furthermore, miRNAs found on the 2α paralogon evolved significantly slower than those on the 2β paralogon.

We next asked if the rate of nucleotide substitution differs significantly across the four paralogons generated from the two vertebrate WGD events (Supp. File 5). Blanc and Wolfe (2004) showed that in *Arabidopsis* the sub-genome with the higher percentage of gene retention exhibits a slower rate of molecular evolution in comparison to the second, gene-poor, sub-genome, and we find the same to with vertebrates, at least with respect to the miRNAs found on sub-genome 2 (Fig. 4B). We aligned the pre-miRNA sequences for each set of miRNA paralogues with at least one paralogue on the α sub-genome and a second on the β sub-genome, and analyzed the concatenated sequences for each of the 13 taxa considered with maximum likelihood (ML, see Materials and Methods). This analysis shows that 2α, the paralogon most enriched for miRNA genes, evolves at a significantly slower rate than 2β, the paralogon most deficient in miRNA genes (*F*(1,48) = 29.43, P<0.0001) (Supp. Table 4). Aligning the entire data set and again analyzing with ML shows nearly an identical set of branch lengths for each of the four paralogons in comparison to the medians calculated from each taxon individually (Fig. 4B; Supp. File 6). Therefore, these data support a model that differences in gene expression, which are correlated to differences in gene mutation, lead to biases in genomic retention following allotetraploidy events.

## DISCUSSION

The Simakov et al. (2020) model for the mode of the vertebrate WGD events was proposed given the disparity between gene retention on the α versus the β paralogons following 2R, but the parity of gene retention between the 1 and 2 sub-genomes following 1R. Further, the timing of these two events was elucidated given that 2R is shared amongst all living gnathostomes, whereas 1R is shared with lampreys. Because MirGeneDB.org explicitly homologizes miRNAs within a taxon and between metazoan taxa, as well as identifies the node of origin of every miRNA locus as well as family, the mode and the timing of the 1R and 2R events can be assessed independently with a non-coding RNA marker. Here, we have shown that miRNAs are retained following WGD events in a manner reflecting the number of WGD events themselves, in both chelicerates and in gnathostome vertebrates (Fig. 2). Further, within gnathostomes, miRNAs follow a similar pattern to the protein-coding repertoire (Fig. 1), with miRNA homeologues enriched on α paralogons relative to β paralogons, but parity seen with ohnologues found on the 1 versus 2 sub-genomes (Fig. 3; Supp.Fig. 2; Table 1). Because the miRNA families with at least one ohnologue are miRNA families also found in cyclostomes, the first genome duplication event happened before the vertebrate LCA. However, miRNA families that are restricted to gnathostomes and are not found in cyclostomes are all found on a single paralogon (Fig. 2, Supp.Fig. 2), confirming that the second WGD occurred after the split between the jawed and jawless fish, but before the gnathostome LCA. Further, given the conservation of mature miRNAs amongst paralogues generated by either the 1R and/or 2R event(s), the biased retention of miRNAs is not due to target interactions with mRNA 3’-UTRs, but instead due to the genomic origin of the miRNA locus itself. Indeed, miRNA paralogues from the α paralogons show both higher expression of miRNA transcripts and slower molecular evolution of the precursor miRNA sequences relative to the β paralogons. Finally, none of the WGDs in either vertebrates or chelicerates resulted in the acquisition of an unusually high number of novel miRNA gene families, and when dramatic increases to miRNA repertoires do occur they are independent of WGD events. Indeed, it is these acquisitions of miRNA families, and not WGDs, which are the likely reason behind increases in morphological and behavioral complexity in metazoans.

### Parity in miRNA Function but Non-Parity in miRNA Retention, Expression and Evolution

An outstanding question concerning WGDs is identifying the mechanism underlying sub-genome dominance following allotetraploidy events. Several hypotheses have been advanced, principally involving either interactions of gene products or expression-level differences between the two newly hybridized genomes. Because homeologues were once orthologues that had independent evolutionary histories before the hybridization event, the co-evolution of interacting gene products in one genome may occur in a different manner relative to the interacting gene products in the second. Thus, similar to a Bateson-Muller-Dobzhansky mechanism of incompatibility (Orr 1996), these two sets of gene products can only work with their partners from the same genome. Therefore, after the hybridization event, one set will be preferentially lost during the rediploidization process relative to the other, even for genes maintained in multiple copies for dosage reasons (Fig. 1B, right). One might expect then that one set of interacting gene products would not necessarily reflect the genomic origin of another, independent set of interacting gene products. However, the fact that both DNA repair proteins *and* transcription factors all show α relative to β dominance (Fig. 1B) suggests that a more likely reason behind sub-genome dominance is the DNA itself, with one set of entire chromosomes preferred over the other.

microRNAs offer an independent test of these ideas. One difficulty in understanding potential incompatibilities between two sets of gene products is simply understanding the detailed nature of the interactions themselves. However, microRNAs interact with messenger RNAs largely through their seven-nucleotide seed sequence (sometimes supplemented with the 3’CR), and thus understanding the potential redundancy between homeologues is, at least in principle, far simpler with miRNAs than with protein sequences. With respect to the gnathostome WGD events, because there are no changes to the seed sequences after either WGD (Supp.Fig. 4), miRNAs from either the α versus the β genome should be interchangeable amongst themselves relative to the genomically preferred mRNA interactors(s). However, not only are miRNAs also strongly preferred on the α sub-genome relative to the β, this preference continues into descendant taxa long after the allotetraploidy event itself (Supp.Fig. 3, Table 2).

Consistent with this continual loss of miRNA paralogues on the β sub-genome relative to the α is the fact that miRNA paralogues are expressed at higher levels on the two α sub-genomes relative to the β sub-genomes (Fig. 4A; Supp. Table 3). Further, the 2α paralogon evolves much slower relative to the 2β paralogon (Fig. 4B; Supp. Table 4), the paralogons with the most and least miRNA paralogues, respectively (Table 2). Indeed, there is a striking and statistically significant relationship between the paralogon placement of miRNAs generated during the 2R events, the expression levels of α versus β paralogues, and the branch lengths leading to the 2α versus the 2β paralogues in 13 representative gnathostome taxa (Fig. 4C). Thus, the gnathostome genome is partitioned into four paralogons, not only in terms of gene content (Simakov et al. 2020; Lamb 2021; Nakatani et al. 2021), but also in terms of miRNA gene expression *and* evolution. How these paralogons maintain their identity for hundreds of millions of years after the 2R events themselves, and if these signals extend to other gene types beyond miRNAs, remain open questions.

### miRNAs, Whole Genome Duplications, and Phenotypic Complexity

WGD events have long enjoyed center stage as the mechanism for driving changes to phenotypic complexity (or species diversity when obvious changes to complexity are not apparent as in teleost fishes relative to other gnathostomes). As originally envisioned by Ohno (1970), because the ancestral vertebrate genome was duplicated in a single event, gene dosage was maintained where needed, with most other ohnologues lost through pseudogenization. However, some of these ohnologues hit upon mutations that gave them new roles to play in vertebrate construction and homeostasis, resulting in a dramatic increase to organismal complexity. Nonetheless, although elegant in its simplicity, this hypothesis is not supported by two observations. First, although *bona fide* instances of sub- and neo-functionalization occurred after both the 2R event and the 3R event (e.g., Prince 2002; Escriva et al. 2006; Kenny et al. 2016; Yamada et al. 2021), the fact that ohno- and homeologues are enriched for gene products whose correct stoichiometric relationships with other gene products is essential, suggests that these instances of sub- and neo-functionalization are likely exaptations (Gould and Vrba 1982), not adaptations, of the WGD events themselves (Freeling and Thomas 2006; Conant et al. 2014; Thompson et al. 2016; Clark and Donoghue 2018). For example, the instances of novelty – whether sub- or neofunctionalization – documented in the *Hox* clusters of mammals and teleosts show that virtually all instances are specific for either the mammal or teleost lineage (Yamada et al. 2021). Thus, like the changes documented herein to mature miRNA sequences (Supp.Fig. 4), these are instances of exaptations that occurred long after the 1R and 2R events. Second, as argued by Donoghue and Purnell (2005), correlations between changes to phenotypic complexity (or diversity) and WGDs are only apparent when extant taxa are considered in isolation. When the fossil record is also considered, this apparent correlation breaks down as there is neither a burst of phenotypic innovation nor a change to diversity that could result from any of the WGDs known to have occurred in vertebrate evolution (see also Clarke et al. 2016; Davesne et al. 2021).

Similar to plants (Clark and Donoghue 2018), the purported relationship between WGDs and complexity is also now not supported by a broader appreciation of the frequency of WGDs throughout metazoans themselves. Indeed, the discovery of multiple WGDs in cyclostomes (Nakatani et al. 2021), in at least two chelicerate lineages (see Fig. 2), as well as the oligochaete worm *Eisenia fetida* (Bhambri et al. 2018), the flatworm *Macrostomum lignano* (Wasik et al. 2015), and even in bdelloid rotifers (Hur et al. 2008; Mark Welch et al. 2008; Flot et al. 2013; Nowell et al. 2018), highlights that there is certainly no necessary correlation between WGD and increases to organismal complexity or species diversity, and certainly no “drive” (Freeling and Thompson 2006) towards increased morphological complexity on the heels of WGDs in either plants or animals. Indeed, as Kenny et al. (2016) emphasized, the fact that *the* classic living fossil – the horseshoe crab – consists of only four extant species and has shown little appreciable change in morphology since the Silurian (Briggs et al. 2012), despite undergoing three WGDs (Fig. 2) sometime before the Early Cretaceous, highlights this absence of correlation, undermining any argument of necessary causality.

To address this disparity between the potential effects WGD had on vertebrate evolution versus chelicerate evolution, Kenny et al. (2016) suggested examining the patterns of miRNA innovation and loss following the WGDs in both. Here, we have revealed three very interesting patterns relevant to their suggestion. First, similar to their principle targets (Fig. 1B, right), miRNA paralogues generated by WGDs are retained after these events in numbers that reflect the number of WGD events themselves (Fig. 2). Nonetheless, this biased retention of miRNAs is not primarily due to neo-functionalizations either in gnathostomes (Supp.Fig. 4) or in the chelicerates (Supp. Fig. 5). Instead, as seen in plants (Abrouk et al. 2012), the retention of miRNA paralogues seems to be driven largely by gene dosage considerations between miRNAs and their target mRNAs, in particular transcription factors (Fig. 1B, right). As several studies have demonstrated, the maintenance of the correct stoichiometry between miRNA mature molecules and the number of miRNA response elements in the 3’-UTRs of mRNAs (Denzler et al. 2014, 2016) is of considerable importance for the normal development and homeostasis of the cell (Broderick and Zamore 2014). Further, within gnathostomes, miRNA paralogues are rarely generated through TGDs, events with the potential to disrupt the stoichiometry between regulator and target. In fact, within the vertebrate set of paralogues generated through the 2R events, not a single shared tandem event occurred over the subsequent 450 million years in any of our 14 representative gnathostomes lineages (Supp.Fig. 3). In fact, just under half (10 of 24 tandem pairs) of these potential, species-specific, gene duplicates have identical pre-miRNA sequences, consistent with the observation that many are likely false positives, the result of mis-assembly of the genome itself (Rhie et al. 2021). Therefore, despite a few potential examples of neo-functionalization (Supp. Figs. 4, 5), these miRNA retention data suggest that very little novelty in terms of regulatory circuits arise following WGDs in either vertebrates or chelicerates.

The second striking pattern is that in all WGD cases examined herein, not once did a WGD event result in a demonstrable increase to the number of miRNA families, only paralogues to previously existing families (Fig. 2) (Heimberg et al. 2008; Fromm et al. 2015). Even with the origin of the vertebrate-specific miRNA repertoire, whose acquisition occurred sometime around the first autotetraploidy event, nonetheless it is likely that most of these new miRNA families were already in place within the pre-1R genome. This is because miRNA families that were certainly in place before the 1R event are distributed in a statistically similar manner (Table 1, *χ*^*2*^ = 1.50, *df*=1, *P*=0.22) to vertebrate-specific singletons that evolved after vertebrates split from urochordates (Supp.Fig. 2). This is a curious observation and raises the question of why WGDs do not generate an influx of new miRNA families given that not only is there a doubling of the number of introns – the predominant source of new miRNAs (Nozawa et al. 2010; Campo-Payssa et al. 2011) – and a doubling of potential target sequences as well. Nonetheless, the absence of miRNA innovation following WGDs in both chelicerates and in vertebrates appears robust.

Instead – and this brings us to the final pattern – pulses of mRNA innovation occur for reasons other than WGDs, reasons that remain speculative at the moment. Nonetheless, it is these dramatic increases in the number of miRNA families and not WGDs that correlate to discrete advents of morphological complexity (Sempere et al. 2006; Heimberg et al. 2008; Peterson et al. 2009; Deline et al. 2018). Three large increases to the rate of miRNA innovation were known across the metazoan kingdom, and in each instance accompanied by an increase in cell-type complexity: at the base of bilaterians, at the base of vertebrates, and at the base of eutherian mammals (Sempere et al. 2006; Hertel et al. 2006; Prochnik et al. 2007; Heimberg et al. 2008; Wheeler et al. 2009; Tarver et al. 2013; Fromm et al. 2015). With each of these documented increased to the miRNA family-level repertoire, clade-specific miRNAs are often expressed in clade-specific tissues (Devor & Peek 2008; Christodoulou et al. 2010; Heimberg et al. 2010; Bartel 2018; Deveale et al. 2021), suggesting that the former might be instrumental in the evolution of the latter (Sempere et al. 2006; Peterson et al. 2009).

A fourth obvious increase to morphological and behavioral complexity occurred in the coleoid cephalopods (i.e., octopus, cuttlefish and squids) after they split from the nautiloid cephalopods, but before the coleoid LCA. Like the horseshoe crab, the octopus, for example, is a striking counter example against the idea that WGDs generate morphological complexity as, despite a notable increase to complexity, there is not a single documented instance of a WGD in this lineage (Albertin et al. 2015). Nonetheless, the octopus has experienced a dramatic increase in the number of miRNA families, with nearly 80 new miRNA families added in this lineage after it split from the Nautilus (Zolotarov et al., in preparation). Unlike placental mammals, coleoids extensively edit their neural mRNA transcriptomes (Albertin et al. 2015; Alon et al. 2015; Liscovitch-Brauer et al. 2017), which may enhance the behavioral repertoire of the coleoid cephalopod. However, the one thing this animal shares with other behaviorally and structurally complex animals like placental mammals is a dramatic influx of novel miRNA families themselves, influxes not caused by WGDs. This might be because with each novel seed sequence added to a genome, a new post-transcriptional regulatory circuit can now be established, bringing additional robustness to the developmental process (Heimberg et al. 2008; Li et al. 2009; Ebert & Sharp 2012; Cassidy et al. 2013; Schmiedel et al. 2015), increasing the heritability of the interaction (Hornstein & Shomron 2006; Peterson et al. 2009; Wu et al. 2009), and ultimately allowing for the evolution of new cell types and functions (Sempere et a. 2006; Deline et al. 2018; Zolotarov et al., in preparation).

## Materials and Methods

All miRNA data, including sequences, expression, and homology assignments, were taken from MirGeneDB.org (https://new.mirgenedb.org/). MirGeneDB identifiers for release 2.1 (Fromm et al., in preparation) were updated from release 2.0 (Fromm et al. 2020) to reflect the paralogon locations of pre-1R miRNA families such that P1 = 1α, P2 = 2α, P3 = 1β, and P4 = 2β (Supp. File 3) except that, as argued by Lamb (2021), the β1 versus β2 of linkage groups B, G, H were switched, as were the α1 versus α2 of linkage group A (paralogon 3 of Lamb 2021). Further, identifies all miRNA paralogue clusters included herein are now numbered so that -P1, for example, is 5’ of -P2, and that all linked genes are given the same linkage identifiers (e.g., Mir-1-P1 and Mir-133-P1 are clustered together, as are Mir-1-P2 and Mir-133-P1). See Fromm et al. (in preparation) for further details and examples.

The miRNA phylogeny (Supp.Fig. 1) was constructed by first aligning the 254 miRNA precursor sequences present in the gnathostome LCA from each of the 32 descendant taxa by eye using both the positions of the RNaseIII cuts as well as the secondary structure of the pre-miRNA molecule as alignment guides. This dataset, consisting of 16,146 nucleotide positions, was analyzed using the CAT-GTR+G (28,000 Cycles) (Supp.Fig. 1) and the GTR +G (2,800 cycles) models in Phylobayes (MPI – version 1.8) with similar results. Convergence was tested using tracecomp and bpcomp (Phylobayes). For the CAT-GTR analyses, we used a burnin of 10,000 cycles and a subsampling frequency of 10. All statistics reported by tracecomp had an effective sample size > 1000 and a relative difference < 0.07 and the bpcomp maxdiff statistic was 0.02, indicating an excellent level of convergence.

To construct an amino acid dataset of equal size to the miRNA dataset, a jackknifing approach was taken. First, we took a set of protein alignments (Braasch et al. 2016) that represents a set of curated orthologous protein families designed for the analysis of vertebrate phylogeny. For each of these protein families, the *Lepisosteus occidentalis* sequence was extracted and the best BLAST [2, 3] hit was found using BLASTp with a maximum e-value of 1e-10 in the proteome of each species present in this study but not that of Braach et al. (2016). Each of these sequences was then blasted back against the *L. occidentalis* proteome and the sequence was added to the orthologous protein family only if it’s best hit was the same protein that was used as the initial query. For the reverse BLAST, as above, the best hit was found using BLASTp with an e-value cutoff of 1e-10. This resulted in 221 gene families for which all species had an ortholog present. These were aligned using mafft 7.429 [4, 5] with default settings and trimmed using TrimAl 1.4 [6] with the -strict option implemented. These trimmed alignments were concatenated to form a superalignment of 80,040 amino acids. From this, five independent jackknife samples were taken using python scripts to randomly select 16,146 sites, which equals the length of the miRNA dataset.

A Bayesian reconstruction of the phylogeny was performed for each dataset using PhyloBayes MPI version 1.8 [7] for between 21,000 and 24,000 cycles under a model of CAT + GTR + 4 discrete gamma categories. Two chains were run for each dataset and, after 2,500 cycles were removed as burn-in, convergence was investigated using bpcomp and tracecomp (part of the PhyloBayes suite). Runs were deemed to have converged if all statistics reported by tracecomp had an effective sample size > 50 and a relative difference < 0.3. After convergence had been reached, a consensus tree was constructed using bpcomp, discarding 2,500 samples as burn-in, from all datasets.

The rate of miRNA nucleotide evolution was done by first aligning pre-miRNA sequences for 170 possible miRNA genes with at least one paralogue on an α paralogon and one paralogue on a β paralogon (Supp. File 4). The resulting alignments for the subset of the 170 genes that could be analyzed for each of the 13 analyzed taxa were assessed using Maximum Likelihood in Paup 4.0a. The GTR model with an estimated rate matrix was used, with a gamma distribution set to 0.5, and the state frequencies empirically derived. A second analysis concatenated each of the 13 taxa into a single super-alignment (Supp. File 5) and was analyzed in exactly the same manner (Fig. 4B).

## Acknowledgements

We would like to thank P. Donoghue and J. Vinther (U. Bristol) for discussion and comments on the manuscript, and the students of Bio63 (Dartmouth College) for their thoughtful suggestions on an earlier draft of this manuscript (in particular B. Bingham, S. Mehta, and C. Shaughnessy). P. Cabot is supported by the Junior Scholars Program (Dartmouth College). OS is supported by the Austrian Science Fund (grant P32190). BF is supported by the Tromsø forskningsstiftelse (TFS) starting grant 20_SG_BF.

---

**Supplemental Table 1.** Paralogue retention in each of the four ancestral paralogons in the gnathostome LCA for genes that encode DNA repair proteins.

| **DNA Repair\*** | **1α** | **2α** | **1β** | **2β** |
| --- | --- | --- | --- | --- |
| Total | 86 | 73 | 12 | 14 |
| 4X | 0 | 0 | 0 | 0 |
| 3X | 0 | 0 | 0 | 0 |
| 2X | 6 | 8 | 2 | 4 |
| 1X | 80 | 65 | 10 | 10 |

\* The difference in parologue retention for genes that encode DNA repair proteins is significantly different between $\alpha$ and $\beta$ paralogs (chi-square = 95.6, df = 1, $P < 0.0001$ ), but not between 1 versus 2 sub-genomes (chi-square = 0.26, $P > 0.6$ ).

**Supplemental Table 2.**
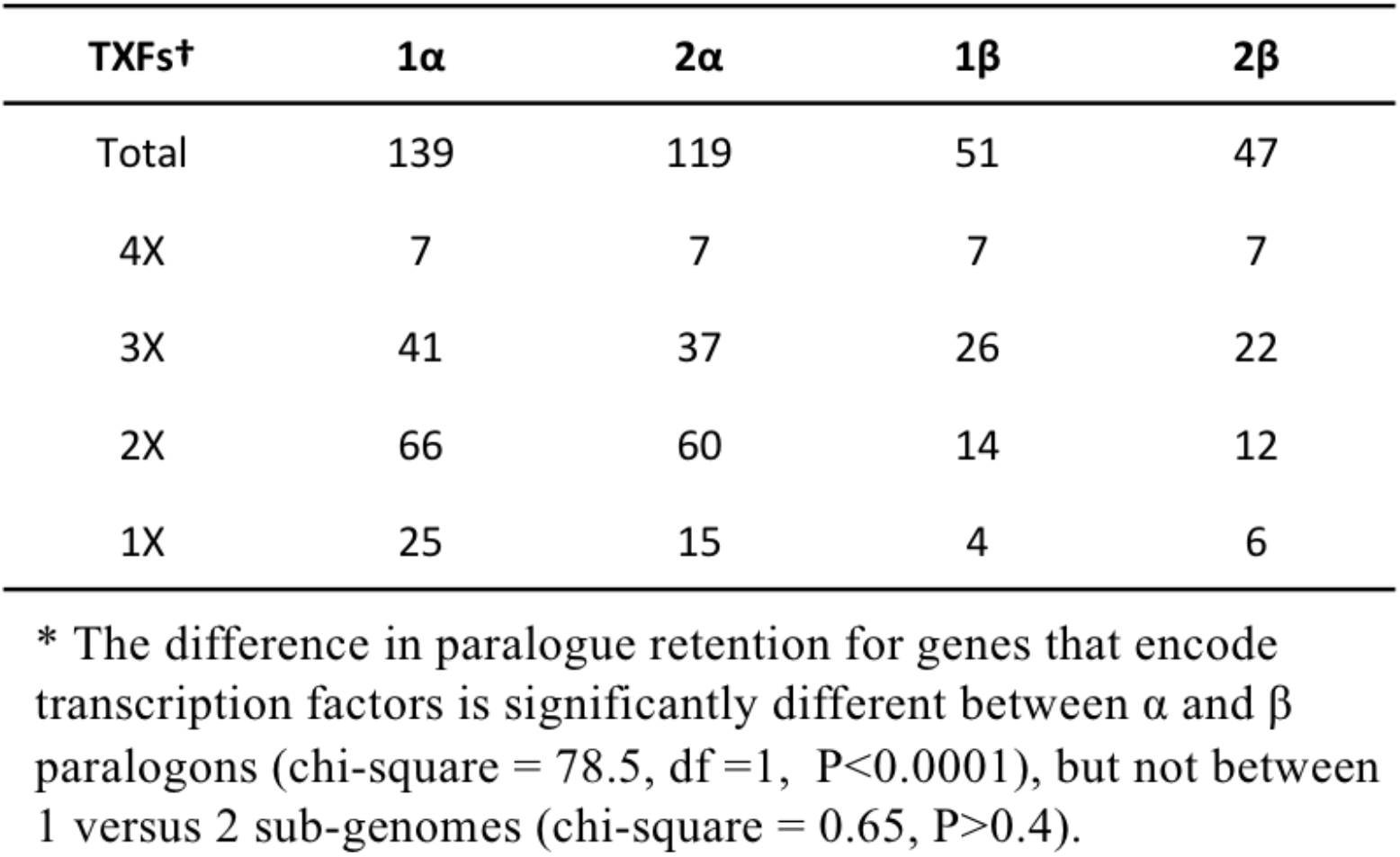
Paralogue retention in each of the four ancestral paralogons in the gnathostome LCA for genes that encode transcription factors.

**Supplemental Table 3.** Median expression values (in reads per million) of unique miRNA mature sequences found on at least one α and one β paralogon.

| Taxon | 1α | 2α | 1β | 2β |
| --- | --- | --- | --- | --- |
| Aca* (40#) | 5,571 | 1,750 | 1,143 | 3,385 |
| Ami (45) | 247 | 242 | 23 | 171 |
| Cmi (46) | 13,833 | 12,339 | 3,955 | 5,513 |
| Cpi (51) | 11,183 | 4,917 | 1,185 | 5,018 |
| Gga (28) | 37,333 | 345,872 | 11,748 | 92,556 |
| Hsa (37) | 39,828 | 69,936 | 28,903 | 59,684 |
| Loc (34) | 10,038 | 9,602 | 4,224 | 5,169 |
| Mdo (31) | 5,576 | 22,675 | 364 | 940 |
| Mun (38) | 1,659 | 852 | 613 | 23 |
| Oan (39) | 2,179 | 31,547 | 1,580 | 1,983 |
| Xtr (38) | 4,761 | 24,886 | 2,523 | 1,218 |
| Median† | 5,574 | 10,971 | 1,383 | 2,684 |

\* Taxon abbreviations: Aca, *Anolis carolinensis*; Ami, *Alligator mississippiensis*; Cmi, *Callorhinchus milii*; Cpi, *Chrysemys picta*; Gga, *Gallus gallus*; Hsa, *Homo sapiens*; Loc, *Lepisosteus oculatus*; Mdo, *Monodelphis domestica*; Mun, *Microcaecilia unicolor*; Oan, *Ornithorhynchus anatinus*; Xtr, *Xenopus tropicalis*.

**Supplemental Table 4.**
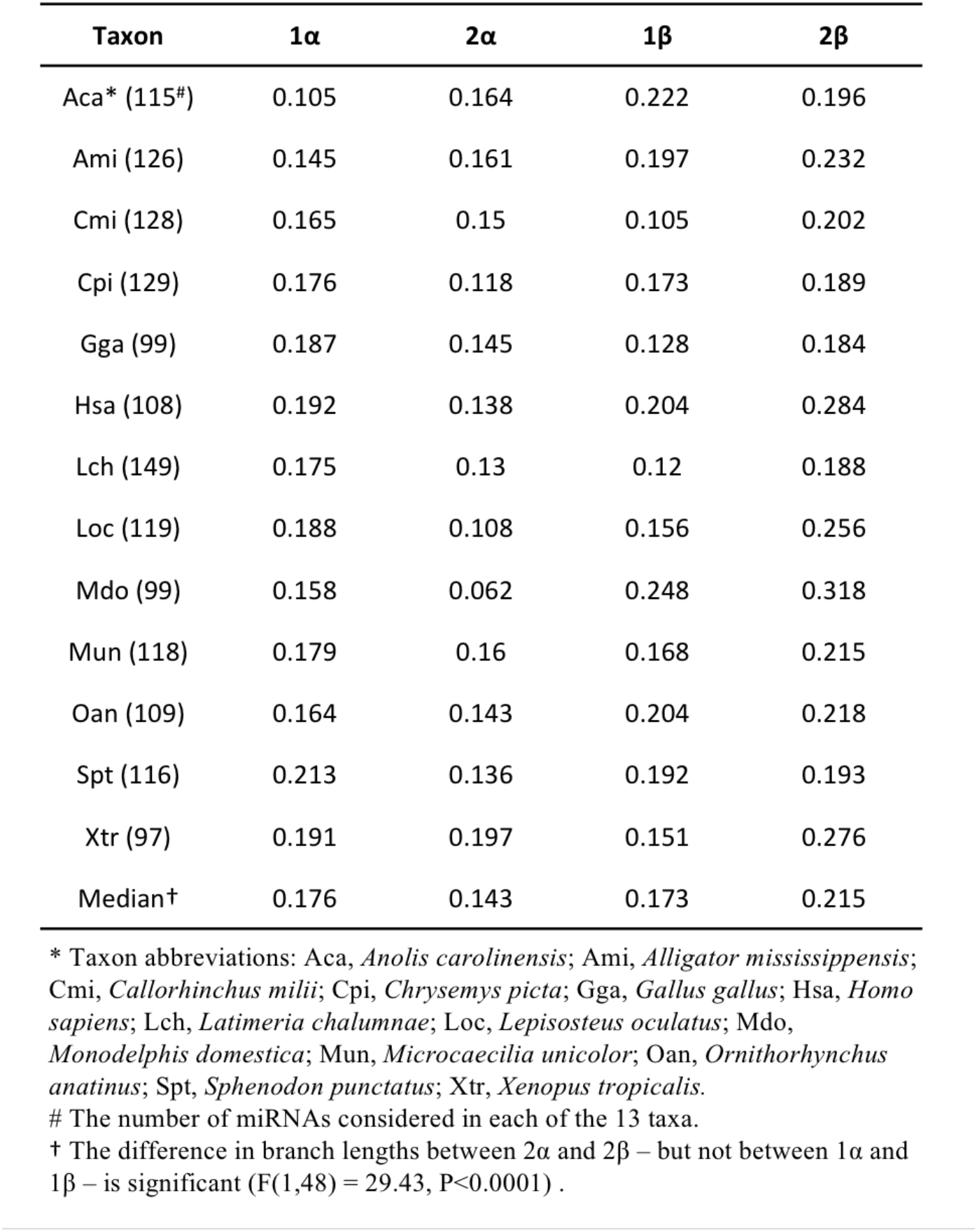
Median branch lengths of pre-miRNA mature sequences found on at least one α and one β paralogon in each of the indicated taxa.

**Supplemental Figure 1.**
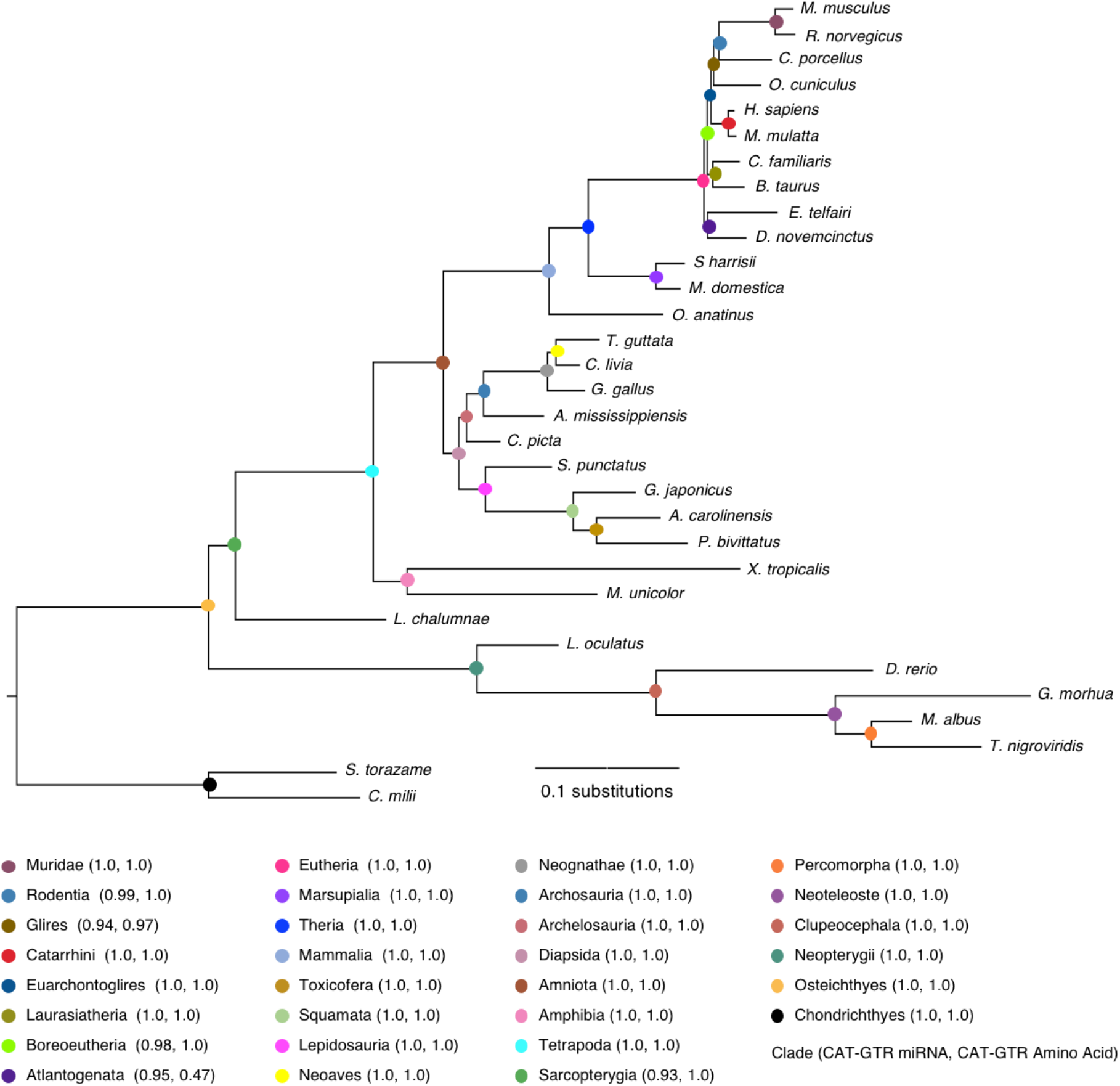
Phylogenetic analysis of 254 pre-miRNA sequences reconstructed as present in the gnathostome LCA (see Mirgenedb.org) from 32 extant taxa (16,146 characters). Each node is color-coded and the support from two different analyses (the posterior probabilities [PP] from CAT-GTR, and GTR, analysis, see materials and methods). Outside of the Boreoeutheria, support values are high, consistent with the orthology assignments of the miRNA loci in MirGeneDB.org.

**Supplemental Figure 2.**
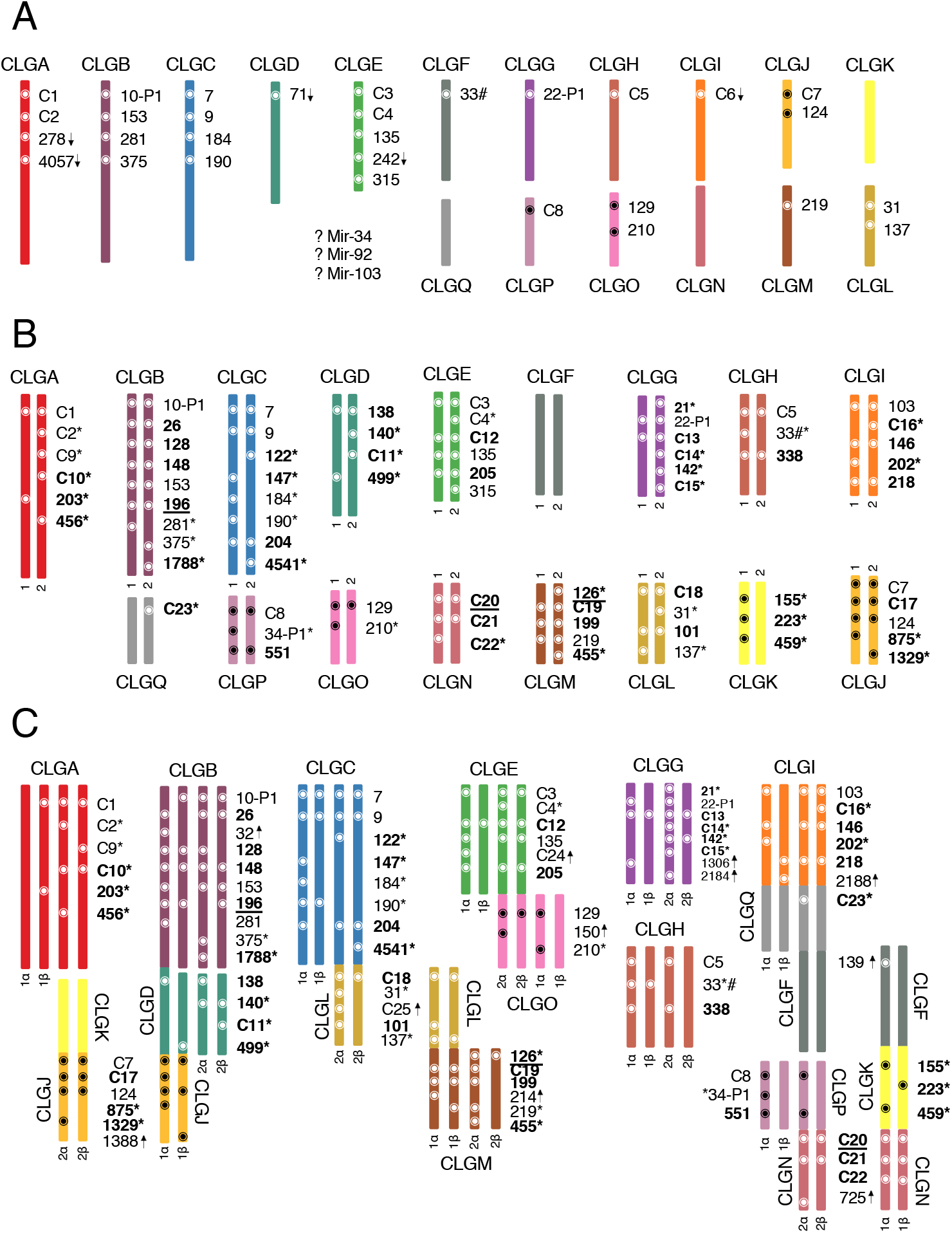
Chromosomal locations of miRNA genes in the last common ancestor of the vertebrate pre-1R (A), vertebrates (B), and gnathostomes. **(C)**. The distribution of miRNA genes are shown across each of the 17 ancestral chordate linkage groups (A-Q). Three ancient miRNA genes or clusters of genes cannot reliably be placed in the vertebrate pre-1R (although they can in the vertebrate LCA). Vertebrate-specific miRNA families are indicated in bold; those in bold underlines arose after the last common ancestor of chordates, but before the last common ancestor of the Olfactores; genes present in the chordate LCA, but lost in the vertebrate LCA, are indicated with a downward arrow; families that evolved after the second genome duplication event are indicated with the upward arrow. The asterisks indicate vertebrate-specific genes present on only one of the two duplicate chromosomes. The pound sign indicates genes that appear to have been displaced from the ancestral linkage group to a different linkage group in the vertebrate LCA. Cluster abbreviations are as follows: C1: Let-7-P1 + Mir-10-P2/P3; C2: 216-P1a/P1b + Mir-217; C3: Mir-29-P1/P2; C4: Mir-96-P1-P3; C5: Mir-193-P1/P2; C6: Mir-252-P1/P2; C7: Mir-1 + Mir-133; C8: Mir-8-P1-P3; C9: Mir-34-P2a/P2b; C10: Mir-192 + Mir-194; C11: Mir-208-P1/P2; C12: Let-7-P2a-P2c; C13: Mir-130-P1-P4; C14: Mir-132-P1/P2; C15: Mir-144 + Mir-451; C16: Mir-143 + Mir-145; C17: Mir-30-P1/P2; C18: Mir-23 + Mir-24 + Mir-27; C19: Mir-181-P1/P2; C20: Mir-15-P1/P2; C21: Mir-17-P1-P4 + Mir-19-P1/P2 + Mir-92-P1/P2; C22: Mir-221-P1/P2; C23: Mir-430-P1-P4; C24: Mir-191 + Mir-425; C25: Mir-34-P3a-d.

**Supplemental Figure 3.**
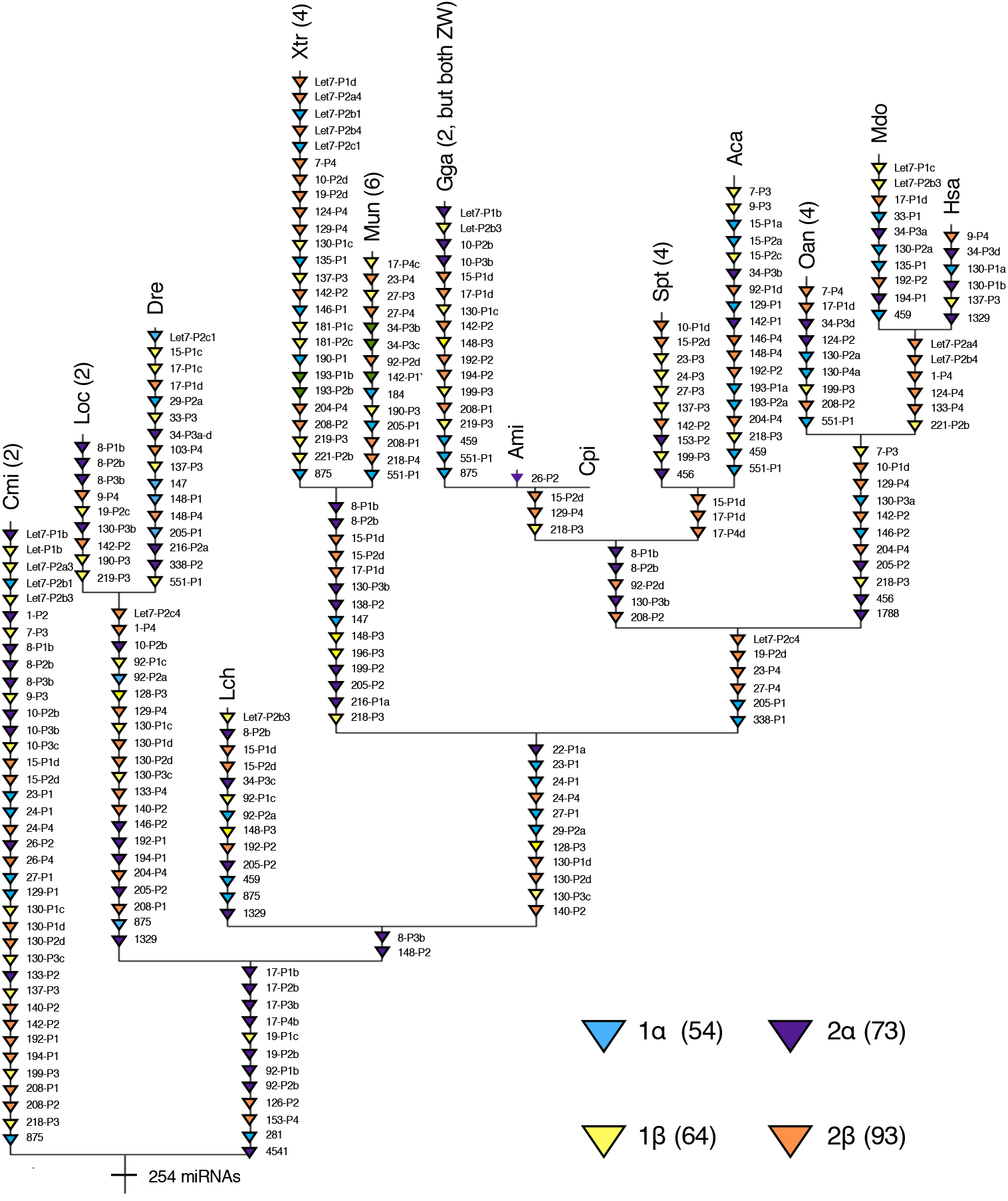
The fate of the 254 miRNAs reconstructed in the gnathostome LCA and their paralogon distribution in 14 extant gnathostome taxa. Despite possessing only about half the miRNA loci (Table 2), miRNAs found on β paralogons are still lost at an enhanced rate relative to α paralogons, a significant difference (*χ*^*2*^ = 32.0, *df* = 3, *P*<0.0001) and one not found with protein-coding loci (Simakov et al. 2020). Numbers in parentheses after the taxon abbreviation are the number of tandem pairs of miRNAs (if any) currently annotated in the latest assembly of that taxon’s genome (see Mirgenedb.org for assembly details for each taxon). No tandem pairs are shared between any two taxa, and most taxa do not have tandem pairs for any of the miRNAs present in the gnathostome LCA. Taxon abbreviations are as follows: Aca, *Anolis carolinensi*; Ami, *Alligator mississippiensis*; Cmi, *Callorhinchus milii* (elephant shark); Cpi, *Chrysemys picta*; Dre, *Danio rerio* (zebrafish); Gga, *Gallus gallus*; Hsa, *Homo sapiens*; Lch, *Latimeria chalumnae*; Loc, *Lepisosteus oculatus* (spotted gar); Mdo, *Monodelphis domestica*; Mun, *Microcaecilia unicolor*; Oan, *Ornithorhynchus anatinus* (platypus); Spi, *Sphenodon punctatus*; Xtr, *Xenopus tropicalis*.

**Supplemental Figure 4.**
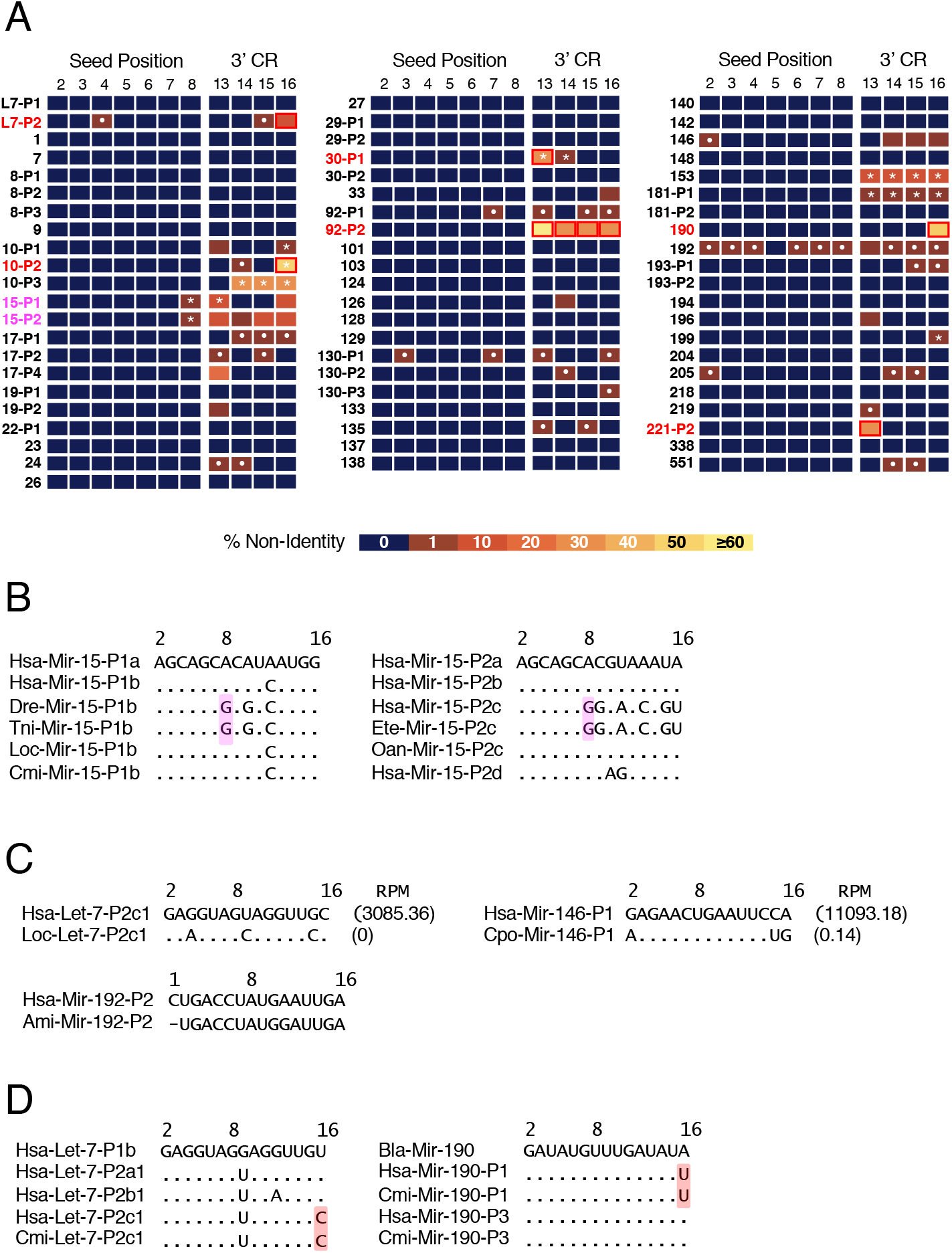
Changes to the seed (nucleotides 2-8) and the 3’-complementary regions (nucleotides 13-16) in miRNA paralogues generated from the gnathostome WGD events. **A**. Sixty-four miRNA genes were present in the vertebrate lineage that had at least two paralogues in the gnathostome lineage resulting from either 1R and/or 2R. Changes to miRNA seed and 3’-complementary regions (CR) were assessed by these regions of the mature miRNA in relation to the consensus sequence for the entire family across the 33 gnathostome taxa present in MirGeneDB v.2.1, and the percentage difference indicated according to the heat map. No changes to the seed sequence – either through seed shifting (Wheeler et al. 2009) or mutations to the seed sequence itself – are the result of these WGD events. Instead, the few changes documented to the seed sequence are the result of mutations that occurred long after 2R. Two of these changes are shared mutations (# symbols) found in specific lineages (magenta), including position 8 in Mir-15-P1 in clupeocephalans or the same position in Mir-15-P2 in eutherian mammals (see panel **B**). Most others though are specific changes that occurred in a single paralogue in a single taxon (white dots), including Let-7-P2c1 in the spotted gar and Mir146-P1 in guinea pig, which might be due to the pseudogenization of the locus (see panel **C**). The only potential instances of neo-functionalization are changes to the 3’ CR regions (red) found in six miRNA paralogues generated by either 1R or 2R (see panel **D**). **B**. The only two examples of shared mutations to the seed sequence are position 8 in Mir-15-P1b in clupeocephalan fish and position 8 of Mir-15-P2c in eutherian mammals (magenta). Because these changes occurred long after 2R they are not examples of neo-functionalization as adaptations of WGD, but, at best, are examples of WGD exaptations. **C**. Some changes to the seed sequence are due to unique changes in a single species due to either due to seed shifting (e.g., Ami-Mir-192-P2) or possibly because of the pseudogenization of the miRNA locus itself (e.g., Loc-Let-7-P2c1 and Cpo-Mir-146-P1) as assessed by the dramatic difference in expression (measured by reads per million [RPM]) between these two potential pseudogenes as compared to their human counterparts. **D**. Potential instances of neo-functionalization are found in the 3’CR of six miRNA paralogues where changes characterize an entire paralogue sub-group relative to other members of the same family (red). For example, both Let-7-P2c1 and Mir-190-P1 have changes in position 16 of the mature miRNA sequence relative to other paralogue subgroups, and/or the single copy member found in invertebrates. These were seen though that resulted from the 2R events (red) including changes to position 16 in both (see panel **E**). Sequence position within the miRNA mature sequence are indicated with the numbers on top; RPM values for Mir-146 are indicated to the right of the sequence. Taxon abbreviations are as follows: Ami, *Alligator mississippiensis*; Bfl, *Branchiostoma floridae* (amphioxus); Cpo, *Cavia porcellus* (guinea pig); Cmi, *Callorhinchus milii* (elephant shark); Dre, *Danio rerio* (zebrafish); Ete, *Echinops telfairi* (tenrec); Hsa, *Homo sapiens*; Loc, *Lepisosteus oculatus* (spotted gar); Oan, *Ornithorhynchus anatinus* (platypus); Tni, *Tetraodon nigroviridis* (pufferfish).

**Supplemental Figure 5.**
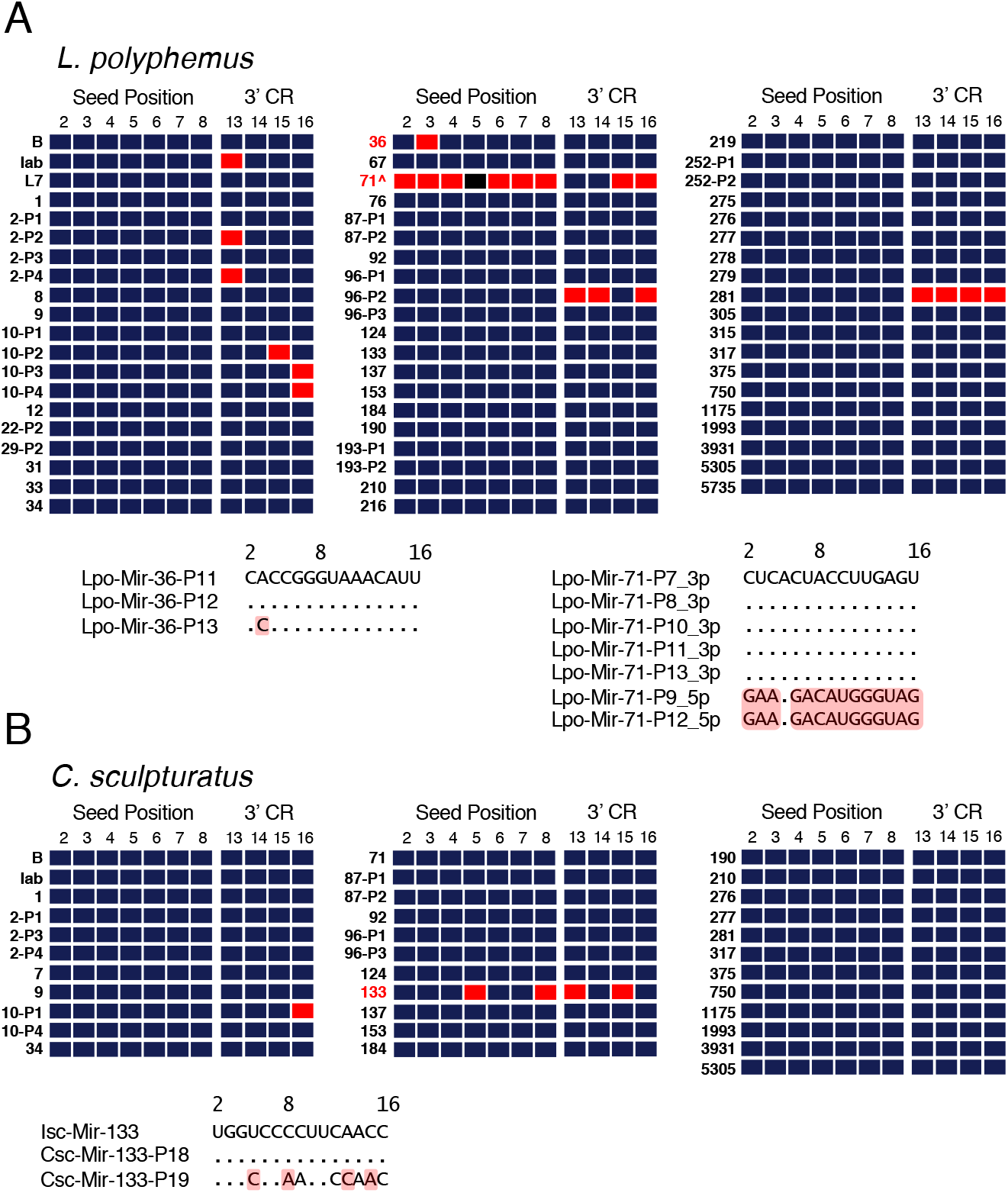
Changes to seed (nucleotides 2-8) and the 3’-complementary regions (nucleotides 13-16) in miRNA paralogues generated from WGDs in chelicerate arthropods. **A**. Cataloguing all changes to the seed and 3’-CR regions in all miRNAs genes present in at least two copies within the horseshoe crab (*L. polyphemus*) genome (red) shows that, like the situation found in gnathostomes (Supp.Fig. 4), most miRNA paralogue sub-groups do not undergo any mutations. Only three instances of potential neo-functionalization are found within the miRNA repertoire of the horseshoe crab, including a change to position 3 in one of the three MIR-36 members, as well as arms switches (Marco et al. 2010; Griffiths-Jones et al. 2011) in two MIR-71 paralogues (^). All other changes are restricted to the 3’ CR regions (red). Note though that because only a single species is represented here, these changes could have occurred long after the three WGD events in the horseshoe crab lineage and thus are not an adaptation, but again an exaptation. **B**. Similarly, cataloguing all changes to the seed and 3’-CR regions in all miRNAs present on two different scaffolds within the scorpion (*C. sculpturatus*) genome (red) shows that, again, most miRNA paralogue sub-groups do not undergo any mutations. This time only a single change to a miRNA mature seed region is seen, *Mir-133-P19*, which has mutation in position 5 of the seed, as well as positions 13 and 15 of the 3’CR. But again, without broader taxon sampling the timing of these changes relative to the single WGD in this lineage remains unknown. Taxon abbreviations: Csc, *Centruroides sculpturatus*; Isc, *Ixodes scapularis* (tick); Lpo, *Limulus polyphemus*.

## References

Abrouk M, Zhang R, Murat F, Li A, Pont C, Mao L, Salse J. 2012. Grass microRNA gene paleohistory unveils new insights into gene dosage balance in subgenome partitioning after whole-genome duplication. Plant Cell 24:1776–1792.

Albertin CB, Simakov O, Mitros T, Wang ZY, Pungor JR, Edsinger-Gonzales E, Brenner S, Ragsdale CW, Rokhsar DS. 2015. The octopus genome and the evolution of cephalopod neural and morphological novelties. Nature 524:220–224.

Alon S, Garrett SC, Levanon EY, Olson S, Graveley BR, Rosenthal JJC, Eisenberg E. 2015. The majority of transcripts in the squid nervous system are extensively recoded by A-to-I RNA editing. elife 4.

Bartel DP. 2009. MicroRNAs: Target recognition and regulatory functions. Cell 136:215–233.

Bartel DP. 2018. Metazoan microRNAs. Cell 173:20–51.

Baskerville S, Bartel DP. 2005. Microarray profiling of microRNAs reveals frequent coexpression with neighboring miRNAs and host genes. RNA 11:241–247.

Berthelot C, Brunet F, Chalopin D, Juanchich A, Bernard M, Noel B, Bento P, Da Silva C, Labadie K, Alberti A, et al. 2014. The rainbow trout genome provides novel insights into evolution after wholegenome duplication in vertebrates. Nat Commun 5:3657–10.

Bhambri A, Dhaunta N, Patel SS, Hardikar M, Bhatt A, Srikakulam N, Shridhar S, Vellarikkal S, Pandey R, Jayarajan R, et al. 2018. Large scale changes in the transcriptome of Eisenia fetida during regeneration. Plos One 13:e0204234.

Birchler JA, Riddle NC, Auger DL, Veitia RA. 2005. Dosage balance in gene regulation: biological implications. Trends. Genet. 21:219–226.

Birchler JA, Veitia RA. 2012. Gene balance hypothesis: connecting issues of dosage sensitivity across biological disciplines. Proc. Natl. Acad. Sci. USA 109:14746–14753.

Blanc G, Wolfe KH. 2004. Functional divergence of duplicated genes formed by polyploidy during Arabidopsis evolution. Plant Cell 16:1679–1691.

Blomme T, Vandepoele K, De Bodt S, Simillion C, Maere S, Van de Peer Y. 2006. The gain and loss of genes during 600 million years of vertebrate evolution. Genome Biol. 7:R43–12.

Braasch I, Gehrke AR, Smith JJ, Kawasaki K, Manousaki T, Pasquier J, Amores A, Desvignes T, Batzel P, Catchen J, et al. 2016. The spotted gar genome illuminates vertebrate evolution and facilitates human-teleost comparisons. Nat. Genet. 48:427–437.

Brancati G, Großhans H. 2018. An interplay of miRNA abundance and target site architecture determines miRNA activity and specificity. Nucleic Acids Res. 46:3259–3269.

Briggs DEG, Siveter DJ, Siveter DJ, Sutton MD, Garwood RJ, Legg D. 2012. Silurian horseshoe crab illuminates the evolution of arthropod limbs. Proc. Natl. Acad. Sci. U.S.A. 109:15702–15705.

Broderick JA, Zamore PD. 2014. Competitive endogenous RNAs cannot alter microRNA function in vivo. Mol. Cell 54:711–713.

Broughton JP, Lovci MT, Huang JL, Yeo GW, Pasquinelli AE. 2016. Pairing beyond the seed supports microRNA targeting specificity. Mol. Cell 64:320–333.

Buggs RJA, Chamala S, Wu W, Tate JA, Schnable PS, Soltis DE, Soltis PS, Barbazuk WB. 2012. Rapid, repeated, and clustered loss of duplicate genes in allopolyploid plant populations of independent origin. Curr. Biol. 22:248–252.

Campo-Paysaa F, Semon M, Cameron RA, Peterson KJ, Schubert M. 2011. microRNA complements in deuterostomes: origin and evolution of microRNAs. Evol Dev 13:15–27.

Cañestro C, Albalat R, Irimia M, Garcia-Fernandez J. 2013. Impact of gene gains, losses and duplication modes on the origin and diversification of vertebrates. Semin. Cell Dev. Biol. 24:83–94.

Cassidy JJ, Jha AR, Posadas DM, Giri R, Venken KJT, Ji J, Jiang H, Bellen HJ, White KP, Carthew RW. 2013. miR-9a minimizes the phenotypic impact of genomic diversity by buffering a transcription factor. Cell 155:1556–1567.

Cheng F, Wu J, Cai X, Liang J, Freeling M, Wang X. 2018. Gene retention, fractionation and subgenome differences in polyploid plants. Nature Plants:1–11.

Christodoulou F, Raible F, Tomer R, Simakov O, Trachana K, Klaus S, Snyman H, Hannon GJ, Bork P, Arendt D. 2010. Ancient animal microRNAs and the evolution of tissue identity. Nature:1–7.

Clark JW, Donoghue PCJ. 2018. Whole-genome duplication and plant macroevolution. Trends Plant Sci. 23:933–945.

Clarke JT, Lloyd GT, Friedman M. 2016. Little evidence for enhanced phenotypic evolution in early teleosts relative to their living fossil sister group. Proc. Natl. Acad. Sci. USA 113:11531–11536.

Comai L. 2005. The advantages and disadvantages of being polyploid. Nat Rev Genet 6:836–846.

Conant GC, Birchler JA, Pires JC. 2014. Dosage, duplication, and diploidization: clarifying the interplay of multiple models for duplicate gene evolution over time. Current Opinion in Plant Biology 19:91–98.

Conant GC, Wolfe KH. 2008. Turning a hobby into a job: how duplicated genes find new functions. Nat Rev Genet 9:938–950.

Davesne D, Friedman M, Schmitt AD, Fernandez V, Carnevale G, Ahlberg PE, Sanchez S, Benson RBJ. 2021. Fossilized cell structures identify an ancient origin for the teleost whole-genome duplication. Proc. Natl. Acad. Sci. USA 118.

Dehal P, Boore J. 2005. Two rounds of whole genome duplication in the ancestral vertebrate. Plos Biol 3:1700–1708.

Deline B, Greenwood JM, Clark JW, Puttick MN, Peterson KJ, Donoghue PCJ. 2018. Evolution of metazoan morphological disparity. Proc. Natl. Acad. Sci. U.S.A. 29:201810575–10.

Denzler R, Agarwal V, Stefano J, Bartel DP, Stoffel M. 2014. Assessing the ceRNA hypothesis with quantitative measurements of miRNA and target abundance. Mol. Cell 54:766–776.

Denzler R, McGeary SE, Title AC, Agarwal V, Bartel DP, Stoffel M. 2016. Impact of MicroRNA Levels, Target-Site Complementarity, and Cooperativity on Competing Endogenous RNA-Regulated Gene Expression. Mol. Cell:1–16.

Desvignes T, Sydes J, Montfort J, Bobe J, Postlethwait JH. 2021. Evolution after whole-genome duplication: Teleost microRNAs. Mol. Biol. Evol. 38:3308–3331.

DeVeale B, Swindlehurst-Chan J, Blelloch R. 2021. The roles of microRNAs in mouse development. Nat Rev Genet:1–17.

Devor EJ, Peek AS. 2008. miRNA Profile of a Triassic Common Mammalian Ancestor and PremiRNA Evolution in the Three Mammalian Reproductive Lineages. TOGENJ 1:22–32.

Dexheimer PJ, Cochella L. 2020. MicroRNAs: From mechanism to organism. Front. Cell Dev. Biol. 8:409.

Donoghue P, Purnell M. 2005. Genome duplication, extinction and vertebrate evolution. Trends Ecol Evol 20:312–319.

Ebert MS, Sharp PA. 2012. Roles for microRNAs in conferring robustness to biological processes. Cell 149:515–524.

Edger PP, McKain MR, Bird KA, VanBuren R. 2018. Subgenome assignment in allopolyploids: challenges and future directions. Current Opinion in Plant Biology 42:76–80.

Edger PP, Pires JC. 2009. Gene and genome duplications: the impact of dosage-sensitivity on the fate of nuclear genes. Chromosome Res 17:699–717.

Escriva H, Bertrand S, Germain P, Robinson-Rechavi M, Umbhauer M, Cartry J, Duffraisse M, Holland L, Gronemeyer H, Laudet V. 2006. Neofunctionalization in vertebrates: the example of retinoic acid receptors. PLoS Genet 2:e102.

Flot J-F, Hespeels B, Li X, Noel B, Arkhipova I, Danchin EGJ, Hejnol A, Henrissat B, Koszul R, Aury J-M, et al. 2013. Genomic evidence for ameiotic evolution in the bdelloid rotifer Adineta vaga. Nature 500:453–457.

Freeling M, Thomas BC. 2006. Gene-balanced duplications, like tetraploidy, provide predictable drive to increase morphological complexity. Genome Res. 16:805–814.

Fromm B, Billipp T, Peck LE, Johansen M, Tarver JE, King BL, Newcomb JM, Sempere LF, Flatmark K, Hovig E, et al. 2015. A uniform system for the annotation of vertebrate microRNA genes and the evolution of the human microRNAome. Annu. Rev. Genet. 49:213–242.

Fromm B, Domanska D, Høye E, Ovchinnikov V, Kang W, Aparicio-Puerta E, Johansen M, Flatmark K, Mathelier A, Hovig E, et al. 2020. MirGeneDB 2.0: the metazoan microRNA complement. Nucleic Acids Res. 48:D132–D141.

Garsmeur O, Schnable JC, Almeida A, Jourda C, D’Hont A, Freeling M. 2013. Two evolutionarily distinct classes of paleopolyploidy. Mol. Biol. Evol. 31:448–454.

Gibson TJ, Spring J. 1998. Genetic redundancy in vertebrates: polyploidy and persistence of genes encoding multidomain proteins. Trends. Genet. 14:46–49.

Glasauer SMK, Neuhauss SCF. 2014. Whole-genome duplication in teleost fishes and its evolutionary consequences. Mol Genet Genomics 289:1045–1060.

Glover NM, Redestig H, Dessimoz C. 2016. Homoeologs: What are they and how do we infer them? Trends Plant Sci. 21:609–621.

Gould SJ, Vrba ES. 1982. Exaptation - a missing term in the science of form. Paleobiology 8:4–15.

Gout J-F, Kahn D, Duret L, Paramecium Post-Genomics Consortium. 2010. The relationship among gene expression, the evolution of gene dosage, and the rate of protein evolution. PLoS Genet 6:e1000944.

Griffiths Jones S, Hui JHL, Marco A, Ronshaugen M. 2011. MicroRNA evolution by arm switching. EMBO Rep 12:172–177.

Heimberg AM, Cowper-Sal-lari R, Semon M, Donoghue PCJ, Peterson KJ. 2010. microRNAs reveal the interrelationships of hagfish, lampreys, and gnathostomes and the nature of the ancestral vertebrate. Proc. Natl. Acad. Sci. USA 107:19379–19383.

Heimberg AM, Sempere LF, Moy VN, Donoghue PCJ, Peterson KJ. 2008. MicroRNAs and the advent of vertebrate morphological complexity. Proc. Natl. Acad. Sci. USA 105:2946–2950.

Hertel J, Lindemeyer M, Missal K, Fried C, Tanzer A, Flamm C, Hofacker IL, Stadler PF, Students of Bioinformatics Computer Labs 2004 and 2005. 2006. The expansion of the metazoan microRNA repertoire. BMC Genomics 7:25–15.

Holland PW, Garcia-Fernandez J, Williams NA, Sidow A. 1994. Gene duplications and the origins of vertebrate development. Dev Suppl:125–133.

Hornstein E, Shomron N. 2006. Canalization of development by microRNAs. Nat. Genet. 38:S20–S24.

Hufton AL, Mathia S, Braun H, Georgi U, Lehrach H, Vingron M, Poustka AJ, Panopoulou G. 2009. Deeply conserved chordate noncoding sequences preserve genome synteny but do not drive gene duplicate retention. Genome Res. 19:2036–2051.

Hughes AL. 1999. Phylogenies of developmentally important proteins do not support the hypothesis of two rounds of genome duplication early in vertebrate history. J Mol Evol 48:565–576.

Huminiecki L, Heldin CH. 2010. 2R and remodeling of vertebrate signal transduction engine. BMC Biol 8:146–21.

Hur JH, Van Doninck K, Mandigo ML, Meselson M. 2009. Degenerate tetraploidy was established before bdelloid rotifer families diverged. Mol. Biol. Evol. 26:375–383.

Kassahn KS, Dang VT, Wilkins SJ, Perkins AC, Ragan MA. 2009. Evolution of gene function and regulatory control after whole-genome duplication: Comparative analyses in vertebrates. Genome Res. 19:1404–1418.

Kenny NJ, Chan KW, Nong W, Qu Z, Maeso I, Yip HY, Chan TF, Kwan HS, Holland PWH, Chu KH, et al. 2016. Ancestral whole-genome duplication in the marine chelicerate horseshoe crabs. Heredity 116:190–199.

Kondrashov FA, Koonin EV. 2004. A common framework for understanding the origin of genetic dominance and evolutionary fates of gene duplications. Trends. Genet. 20:287–290.

Lamb TD. 2021. Analysis of paralogons, origin of the vertebrate karyotype, and ancient chromosomes retained in extant species. Genome Biology and Evolution 13.

Lambert SA, Jolma A, Campitelli LF, Das PK, Yin Y, Albu M, Chen X, Taipale J, Hughes TR, Weirauch MT. 2018. The human transcription factors. Cell 172:650–665.

Leite DJ, Ninova M, Hilbrant M, Arif S, Griffiths-Jones S, Ronshaugen M, Mcgregor AP. 2016. Pervasive microRNA duplication in chelicerates: Insights from the embryonic microRNA repertoire of the spider Parasteatoda tepidariorum. Genome Biol. 8:2133–2144.

Li X, Cassidy JJ, Reinke CA, Fischboeck S, Carthew RW. 2009. A microRNA imparts robustness against environmental fluctuation during development. Cell 137:273–282.

Liscovitch-Brauer N, Alon S, Porath HT, Elstein B, Unger R, Ziv T, Admon A, Levanon EY, Rosenthal JJC, Eisenberg E. 2017. Trade-off between Transcriptome Plasticity and Genome Evolution in Cephalopods. Cell 169:191–202.e11.

Lundin L-G, Larhammar D, Hallböök F. 2003. Numerous groups of chromosomal regional paralogies strongly indicate two genome doublings at the root of the vertebrates. J Struct Funct Genomics 3:53–63.

Makino T, Hokamp K, McLysaght A. 2009. The complex relationship of gene duplication and essentiality. Trends. Genet. 25:152–155.

Makino T, McLysaght A. 2010. Ohnologs in the human genome are dosage balanced and frequently associated with disease. Proc. Natl. Acad. Sci. U.S.A. 107:9270–9274.

Marco A, Hui JHL, Ronshaugen M, Griffiths-Jones S. 2010. Functional shifts in insect microRNA evolution. Genome Biol.

Mark Welch DB, Welch JLM, Meselson M. 2008. Evidence for degenerate tetraploidy in bdelloid rotifers. Proc. Natl. Acad. Sci. U.S.A. 105:5145–5149.

Nakanishi K. 2016. Anatomy of RISC: how do small RNAs and chaperones activate Argonaute proteins? WIREs RNA 7:637–660.

Nakatani Y, Shingate P, Ravi V, Pillai NE, Prasad A, McLysaght A, Venkatesh B. 2021. Reconstruction of proto-vertebrate, proto-cyclostome and proto-gnathostome genomes provides new insights into early vertebrate evolution. Nat Commun:1–14.

Nong W, Qu Z, Li Y, Barton-Owen T, Wong AYP, Yip HY, Lee HT, Narayana S, Baril T, Swale T, et al. 2021. Horseshoe crab genomes reveal the evolution of genes and microRNAs after three rounds of whole genome duplication. Communications Biology:1–11.

Nowell RW, Almeida P, Wilson CG, Smith TP, Fontaneto D, Crisp A, Micklem G, Tunnacliffe A, Boschetti C, Barraclough TG. 2018. Comparative genomics of bdelloid rotifers: Insights from desiccating and nondesiccating species. Plos Biol 16:e2004830.

Nozawa M, Miura S, Nei M. 2010. Origins and evolution of microRNA genes in Drosophila species. Genome Biol. 2:180–189.

Ohno S. 1970. Evolution by gene duplication. Berlin, New York, Springer-Verlag

Orr HA. 1996. Dobzhansky, Bateson, and the genetics of speciation. Genetics 144:1331–1335.

Papp B, Pal C, Hurst LD. 2003. Dosage sensitivity and the evolution of gene families in yeast. Nature 424:194–197.

Peterson KJ, Dietrich MR, McPeek MA. 2009. MicroRNAs and metazoan macroevolution: insights into canalization, complexity, and the Cambrian explosion. Bioessays 31:736–747.

Prince V. 2002. The Hox Paradox: More complex(es) than imagined. Dev. Biol. 249:1–15.

Prochnik SE, Rokhsar DS, Aboobaker AA. 2007. Evidence for a microRNA expansion in the bilaterian ancestor. Dev Genes Evol 217:73–77.

Putnam NH, Butts T, Ferrier DEK, Furlong RF, Hellsten U, Kawashima T, Robinson-Rechavi M, Shoguchi E, Terry A, Yu J-K, et al. 2008. The amphioxus genome and the evolution of the chordate karyotype. Nature 453:1064–U3.

Rhie A, McCarthy SA, Fedrigo O, Damas J, Formenti G, Koren S, Uliano-Silva M, Chow W, Fungtammasan A, Kim J, et al. 2021. Towards complete and error-free genome assemblies of all vertebrate species. Nature:1–32.

Sacerdot C, Louis A, Bon C, Berthelot C, Crollius HR. 2018. Chromosome evolution at the origin of the ancestral vertebrate genome. Genome Biol.:1–15.

Schirle NT, Sheu-Gruttadauria J, MacRae IJ. 2014. Structural basis for microRNA targeting. Science 346:608–613.

Schmiedel JM, Klemm SL, Zheng Y, Sahay A, Blüthgen N, Marks DS, van Oudenaarden A. 2015. MicroRNA control of protein expression noise. Science 348:128–132.

Schnable JC, Springer NM, Freeling M. 2011. Differentiation of the maize subgenomes by genome dominance and both ancient and ongoing gene loss. Proc. Natl. Acad. Sci. U.S.A. 108:4069–4074.

Schwager EE, Sharma PP, Clarke T, Leite DJ, Wierschin T, Pechmann M, Akiyama-Oda Y, Esposito L, Bechsgaard J, Bilde T, et al. 2017. The house spider genome reveals an ancient whole-genome duplication during arachnid evolution. :1–27.

Sempere LF, Cole CN, McPeek MA, Peterson KJ. 2006. The phylogenetic distribution of metazoan microRNAs: Insights into evolutionary complexity and constraint. J. Exp. Zool. 306B:575–588.

Seoighe C, Gehring C. 2004. Genome duplication led to highly selective expansion of the Arabidopsis thaliana proteome. Trends. Genet. 20:461–464.

Session AM, Uno Y, Kwon T, Chapman JA, Toyoda A, Takahashi S, Fukui A, Hikosaka A, Suzuki A, Kondo M, et al. 2016. Genome evolution in the allotetraploid frog Xenopus laevis. Nature 538:336–343.

Shingate P, Ravi V, Prasad A, Tay B-H, Garg KM, Chattopadhyay B, Yap L-M, Rheindt FE, Venkatesh B. 2020. Chromosome-level assembly of the horseshoe crab genome provides insights into its genome evolution. Nat Commun:1–13.

Shingate P, Ravi V, Prasad A, Tay B-H, Venkatesh B. 2020. Chromosome-level genome assembly of the coastal horseshoe crab (Tachypleus gigas). Mol Ecol Resour 20:1748–1760.

Sidow A. 1996. Gen(om)e duplications in the evolution of early vertebrates. Curr Opin Genet Dev 6:715–722.

Simakov O, Marlétaz F, Yue J-X, O’Connell B, Jenkins J, Brandt A, Calef R, Tung C-H, Huang T-K, Schmutz J, et al. 2020. Deeply conserved synteny resolves early events in vertebrate evolution. Nat. Ecol. Evol.: 1–22.

Singh PP, Affeldt S, Cascone I, Selimoglu R, Camonis J, Isambert H. 2012. On the expansion of “dangerous” gene repertoires by whole-genome duplications in early vertebrates. Cell Rep. 2:1387–1398.

Tarver JE, Sperling EA, Nailor A, Heimberg AM, Robinson JM, King BL, Pisani D, Donoghue PCJ, Peterson KJ. 2013. miRNAs: small genes with big potential in metazoan phylogenetics. Mol. Biol. Evol. 30:2369–2382.

Thomas BC, Pedersen B, Freeling M. 2006. Following tetraploidy in an Arabidopsis ancestor, genes were removed preferentially from one homeolog leaving clusters enriched in dose-sensitive genes. Genome Res. 16:934–946.

Thompson A, Zakon HH, Kirkpatrick M. 2016. Compensatory drift and the evolutionary dynamics of dosage-sensitive duplicate genes. Genetics 202:765–774.

Van de Peer Y, Maere S, Meyer A. 2009. The evolutionary significance of ancient genome duplications. Nature:1–8.

Van de Peer Y, Mizrachi E, Marchal K. 2017. The evolutionary significance of polyploidy. Nat Rev Genet 18:411–424.

Veitia RA. 2002. Exploring the etiology of haploinsufficiency. Bioessays 24:175–184.

Veitia RA. 2003. Nonlinear effects in macromolecular assembly and dosage sensitivity. Journal of Theoretical Biology 220:19–25.

Veitia RA. 2009. A generalized model of gene dosage and dominant negative effects in macromolecular complexes. The FASEB Journal 24:994–1002.

Veron AS, Kaufmann K, Bornberg-Bauer E. 2007. Evidence of interaction network evolution by wholegenome duplications: a case study in MADS-box proteins. Mol. Biol. Evol. 24:670–678.

Wasik K, Gurtowski J, Zhou X, Ramos OM, Delás MJ, Battistoni G, Demerdash El O, Falciatori I, Vizoso DB, Smith AD, et al. 2015. Genome and transcriptome of the regeneration-competent flatworm, Macrostomum lignano. Proc. Natl. Acad. Sci. USA 112:12462–12467.

Wee LM, Flores-Jasso CF, Salomon WE, Zamore PD. 2012. Argonaute Divides Its RNA Guide into Domains with Distinct Functions and RNA-Binding Properties. Cell 151:1055–1067.

Wheeler BM, Heimberg AM, Moy VN, Sperling EA, Holstein TW, Heber S, Peterson KJ. 2009. The deep evolution of metazoan microRNAs. Evol Dev 11:50–68.

Wu C-I, Shen Y, Tang T. 2009. Evolution under canalization and the dual roles of microRNAs: a hypothesis. Genome Res. 19:734–743.

Xu P, Xu J, Liu G, Chen L, Zhou Z, Peng W, Jiang Y, Zhao Z, Jia Z, Sun Y, et al. 2019. The allotetraploid origin and asymmetrical genome evolution of the common carp Cyprinus carpio. Nat Commun:1–11.

Yamada K, Maeno A, Araki S, Kikuchi M, Suzuki M, Ishizaka M, Satoh K, Akama K, Kawabe Y, Suzuki K, et al. 2021. An atlas of seven zebrafish hox cluster mutants provides insights into sub/neofunctionalization of vertebrate Hox clusters. Development 148.

